# Divergent effects of 3-nitrooxypropanol on enteric methane emissions in Holstein and Brown Swiss cows, and its lack of synergy with *Acacia mearnsii* tannin extract

**DOI:** 10.1101/2024.11.24.625036

**Authors:** M. Z. Islam, S. E. Räisänen, T. He, C. Kunz, Y. Li, X. Ma, A. M. Serviento, K. Wang, M. Wang, Z. Zeng, M. Niu

## Abstract

The objectives of this study were to investigate the combined effects of 3-nitrooxypropanol (**3-NOP**) and *Acacia mearnsii* tannin extract (**TAN**), and their interactions with dairy cattle breed [Brown Swiss (**BS**) vs. Holstein Friesian (**HF**)], on lactational performance, and enteric methane (**CH_4_**) emissions. Sixteen multiparous mid-lactation cows, including 8 BS and 8 HF cows were used in a split-plot design, with breed as the main plot. Cows within each subplot were arranged in a replicated 4 × 4 Latin Square design with a 2 × 2 factorial arrangement of treatments across four 24-d periods. The experimental diets were: 1) **CON** (basal total mixed ration), 2) 3-NOP (60 mg/kg DM), 3) TAN (3% of DM), and 4) 3-NOP + TAN. Spot samples of urine, feces, and gas emissions (via GreenFeed) were collected at the end of each period 8 times over 3 days. No 3-NOP × TAN × Breed interactions were observed for DM intake (**DMI**), milk production, or enteric gas emissions, except for CH_4_ yield (g/kg DMI) and CO_2_ production. Breed influenced DMI, milk production, and component yields, with HF cows consuming 3.7 kg/d more DMI, producing 9.3 kg/d more milk, and achieving greater feed efficiency and higher milk component yields than BS cows. Milk yield and energy-corrected milk (**ECM**) tended to increase in HF but tended to decrease in BS cows by 3-NOP. Cows fed TAN had 1 kg/d lower DMI with the tendency for 3-NOP × TAN showed greater reduction when TAN was fed alone, but milk yield, ECM, and feed efficiency remained unchanged. Cows fed TAN exhibited 18% lower milk urea nitrogen (**N**) concentration and 23.0% lower urinary N but 36.7% greater fecal N excretions as a percentage of daily N intake. A 3-NOP × Breed interaction was observed in CH_4_ production (g/d), with a 21.7% reduction in HF, and a 13.0% reduction in BS. Similarly, there were 3-NOP × Breed tendencies in CH_4_ yield and intensity (g/kg ECM), with reductions in HF cows of 21.8% and 23.4%, respectively, compared to 11.0% and 10.8% in BS cows. In conclusion, there were no synergistic or additive effects between 3-NOP and TAN on enteric CH_4_ mitigation. The enteric CH_4_ emission mitigating effect of 3-NOP was more pronounced in HF cows than in BS cows. Further research is needed to understand breed-specific responses and to optimize CH_4_ mitigation strategies for inclusion in national greenhouse gas inventories.

**Implications:** This study demonstrated that the feed additive 3-nitrooxypropanol reduces enteric methane emissions in dairy cows, with a greater reduction observed in Holsteins (22%) compared to Brown Swiss (13%) cows. While combining 3-NOP with *Acacia mearnsii* tannin extract did not further reduce methane. Feeding *Acacia mearnsii* extract decreased nitrogen excretion, potentially reducing environmental nitrogen pollution from manure. Neither additive substantially impacted milk production; however, 3-nitrooxypropanol tended to increase milk yield in Holsteins while reducing it in Brown Swiss. Overall, this study suggests that enteric methane mitigation efficacy of dietary strategies for different cattle breeds should be further investigated.

## Introduction

In dairy cattle production systems, enteric methane (**CH_4_**) constitutes the principal source of greenhouse gas (**GHG**) emissions, accounting for more than 70% of the total on-farm non-carbon dioxide emissions (Eckard and Clark, 2018). Mitigating enteric CH_4_ emissions is crucial in the dairy cattle industry because, despite being a short-lived flow gas that remains stagnant due to its simultaneous emission and destruction rates, it has a significantly higher global warming potential than other GHG (Allen et al., 2018).

To mitigate enteric CH_4_ emissions, strategies such as dietary interventions, management improvements, enhancing forage quality, and rumen manipulation have been proposed (Hristov et al., 2013). Their effectiveness, especially in altering rumen metabolism through feed additives, has been thoroughly reviewed by Honan et al. (2021) and investigated in a meta-analysis by Arndt et al. (2022). Tannins, a group of polyphenolic plant secondary compounds, are recognized for their protein-binding ability and potential to impact ruminal fermentation, diet digestibility, feed intake, production performance, and enteric CH_4_ emissions (Jayanegara et al., 2012; Aboagye et al., 2018; Herremans et al., 2020). *Acacia mearnsii* (**TAN**, or Black wattle) produces a condensed tannin-rich extract from its bark (Ahmed et al., 2005). Tannins reduce CH_4_ emissions by binding to nutrients, making them inaccessible to microbes, which impairs ruminal fiber degradation (Frutos et al., 2004) and exerts bacteriostatic effects against methanogens (Bodas et al., 2012). However, their efficacy varies based on the type, condensed or hydrolyzable tannin, and on plant maturity, which influences tannin concentrations and makes their impact on CH_4_ reduction highly variable (Aboagye and Beauchemin, 2019). A dose over 5% of DM can also reduce feed intake, limiting their practical use in CH_4_ mitigation (Jayanegara et al., 2012). While some studies note a consistent CH_4_ yield reduction with higher tannin levels, the optimal inclusion rate is still undetermined. For instance, a meta-analysis showed that an average inclusion rate of tannin at 1.5% of DM lowered CH_4_ production and yield by 10% and 5.9%, respectively, without impacting growth or feed efficiency in beef cattle (Orzuna-Orzuna et al., 2021).

A potent CH_4_ inhibitor 3-nitrooxypropanol (**3-NOP**; Bovaer®; DSM Nutrition Products Ltd., Switzerland), has recently been approved by the European Food Safety Authority (EFSA, 2021) and the Food and Drug Administration (FDA, 2024) for use in enteric CH_4_ mitigation in dairy cows, highlighting its potential for global adoption. It targets the enzyme methyl-coenzyme M reductase (**MCR**) to impede the final step of methanogenesis (Duin et al., 2016). Studies indicate that 3-NOP can reduce CH_4_ yield by 26–54%, with its effectiveness varying based on dosage and the nutrient profile of the diet, including different levels of NDF, fat, and starch (Dijkstra et al., 2018; Yu et al., 2021; Kebreab et al., 2023). Notably, a meta-analysis by Kebreab et al. (2023) estimated that a 1% decrease in dietary NDF may enhance the CH_4_ reduction efficacy of 3-NOP by 0.915%, suggesting that high-fiber diets diminish its mitigation efficacy. In our recent study by Ma et al. (2024) with a high-NDF diet (43% of DM), we observed an unexpected variation in the efficacy of 3-NOP between Holstein Friesian (**HF**) and Brown Swiss (**BS**) cows by 18 and 8% reduction in CH_4_ production, respectively. As such an effect has not been observed in previous studies, this warrants further investigation into breed and 3-NOP interactions to optimize its use across different ruminant populations.

The individual effects of 3-NOP and TAN on reducing enteric CH_4_ emissions in dairy cattle have been well-documented. However, the potential for increased efficiency through their combined effect on ruminal CH_4_ emissions has not been investigated previously. Therefore, the objectives of this study were (i) to investigate the combined effects (synergistic or additive) of supplementing 3-NOP and TAN to potentially mitigate enteric CH_4_ emissions, and (ii) to evaluate the possible interactions between breeds of cows with 3-NOP or TAN. We hypothesized that the distinct mode of action of 3-NOP (targeting MCR) and TAN (impairing fiber degradation and acting bacteriostatically against methanogens) would inhibit methanogenesis and alter the rumen ecosystem and their combined supplementation will result in synergistic and/or additive effects on CH_4_ inhibition. Furthermore, we anticipated that the magnitude and nature of these effects will vary between cow breeds, due to inherent differences in rumen microbial fermentation and how these breeds process and respond to feed additives.

## Material and methods

The experiment was conducted at ETH Zürich, AgroVet-Strickhof research station (Lindau, Zurich, Switzerland) from March to June 2023.

### Animals, experimental design, and diets

Sixteen multiparous lactating dairy cows were arranged in a split-plot design, where the main plot was the breed of cows (8 HF and 8 BS cows). Statistical power calculations were conducted in R, using the methodology outlined by Stroup (1999), which confirmed that the power (> 0.8) was sufficient for detecting the interaction between 3-NOP and breed in the current design and sample size. Given the previously observed enteric CH_4_ mitigation efficacy of 3-NOP in BS cows (8%) and HF cows (18%; Ma et al., 2024), the effect size used for power calculation was 10% for enteric CH_4_ production. The HF cows averaged (mean ± SD) 134 ± 31 days-in-milk (**DIM**), 35.8 ± 6.04 kg/d milk yield, and 710 ± 68.0 kg BW, and the BS averaged 161 ± 40 DIM, 27.0 ± 6.24 kg/d milk yield and 710 ± 63.0 kg BW at the start of the experiment. Within each subplot, cows were randomly assigned to treatment sequences in a replicated 4 × 4 Latin Square design with a 2 × 2 factorial arrangement of 2, 3-NOP supplementation categories [Yes/No] and 2 TAN categories [Yes/No; Table 1]. The experiment had 4, 24-d periods, of which 17 d of dietary adaptation and 7 d of sampling. The 4 dietary treatments were: 1) the basal control diet (**CON**), 2) CON with 3-NOP supplementation (60 mg/kg DM, provided through Bovaer®10 at 600 mg/kg DM), 3) CON with tannin extract from *Acacia mearnsii* (TAN; 3% of DM, extract containing 51.8% total tannin per kg of DM and condensed (**CT**): hydrolyzable tannin (**HT**) ratio was 0.219; Weibull Black, TENAC, Montenegro, Brazil), and 4) 3-NOP + TAN. A 3% DM dose of TAN was chosen to balance CH_4_ reduction while maintaining DM intake (**DMI**) and production performance (Denninger et al., 2020), as higher doses (5% of DM; Jayanegara et al., 2012), could negatively impact feed intake and production. The experimental diet ingredients and chemical composition are presented in Table 1. Diets were provided *ad libitum* at 110% of the intake from the previous day. The cows were fed twice daily at 0800 and 1800 h, at a proportion of 60 and 40%, respectively. Diets without 3-NOP received a carrier substance of silicon dioxide and 1, 2-propanediol in a similar proportion to 3-NOP treatment. For incorporation into the basal total mixed ration (**TMR**), we mixed either 3-NOP or a placebo with a ground corn base (65% corn, 30% Bovaer® 10, and 5% sunflower oil) to achieve a target concentration of 60 mg/kg DM at each feeding while TAN and 3-NOP+TAN were added after mixing with the balanced amount of ground corn. The target concentration was achieved by incorporating 2% premix ground corn carrier, which contained 30 g of 3-NOP/kg (as-fed basis). The prepared premixes were top-dressed on the basal diet and uniformly mixed using metal forks in the feed bin (free-stall) and feeding plates (tie-stall) for individual cows. This method ensured thorough blending of the premixes with the TMR at each feeding. During the first 17-d of diet adaptation, the cows were housed in a free-stall barn with free access to drinking water. Individual feed intake was consistently monitored throughout the study using troughs equipped with electronic load cells (Waagen Döhrn GmbH, Germany), and an identification system (American Calan Inc., Northwood, NH, USA), each assigned to one cow. From 18 to 24-d of the experiment, the cows were moved to a tie-stall barn with a feed monitoring system using individual scales (PFA575 Mettler Toledo, Greifensee, Switzerland). Fresh drinking water and feed were provided ad libitum, and cow comfort was ensured with a rubber-matted cubicle bedded with straw and wood shavings. At the end of each of the 4 periods, the BW of the cows was recorded using a TERRA ET scale (n = 63; Waagen Döhrn, Germany).

**Table 1.**
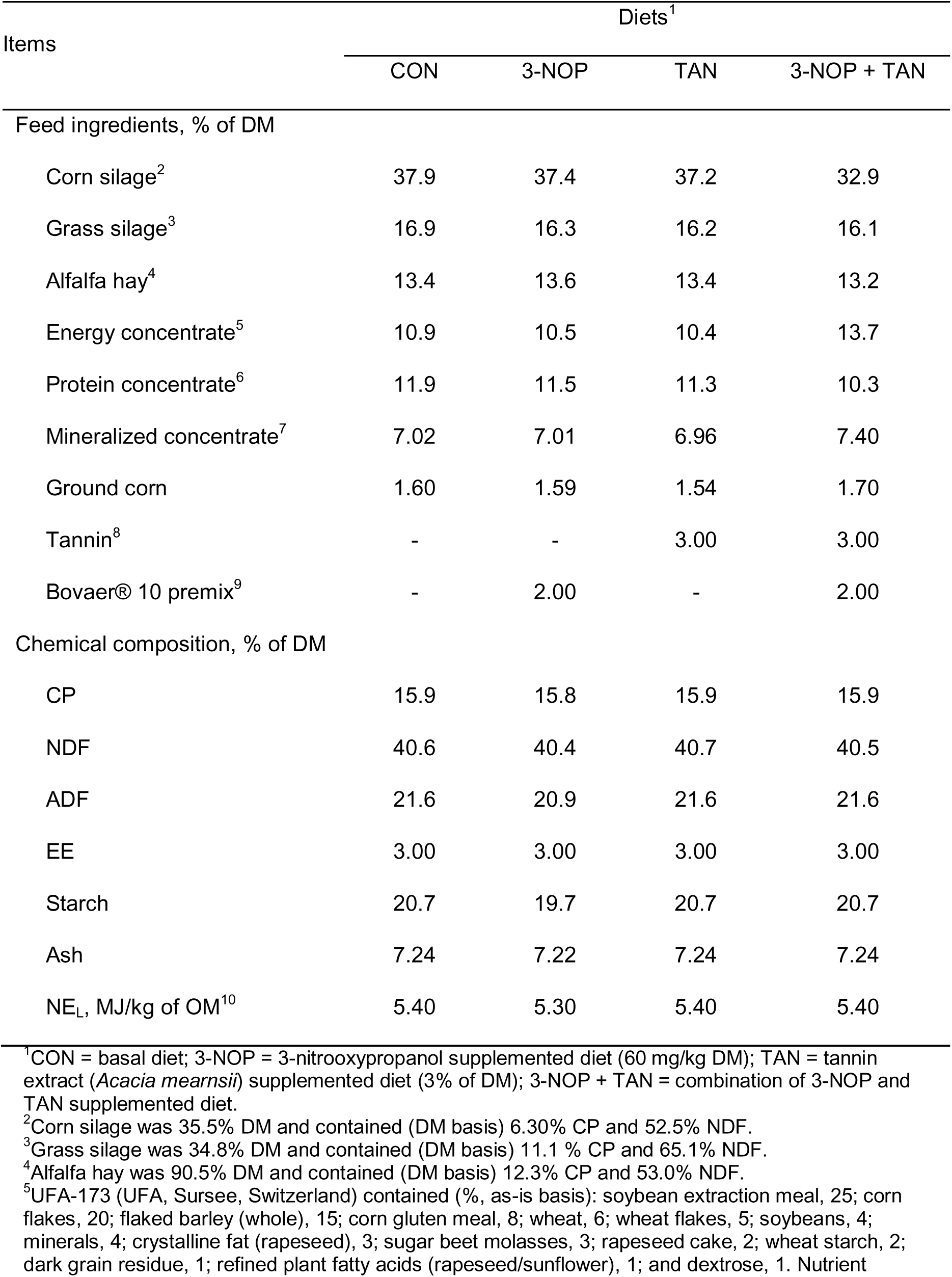
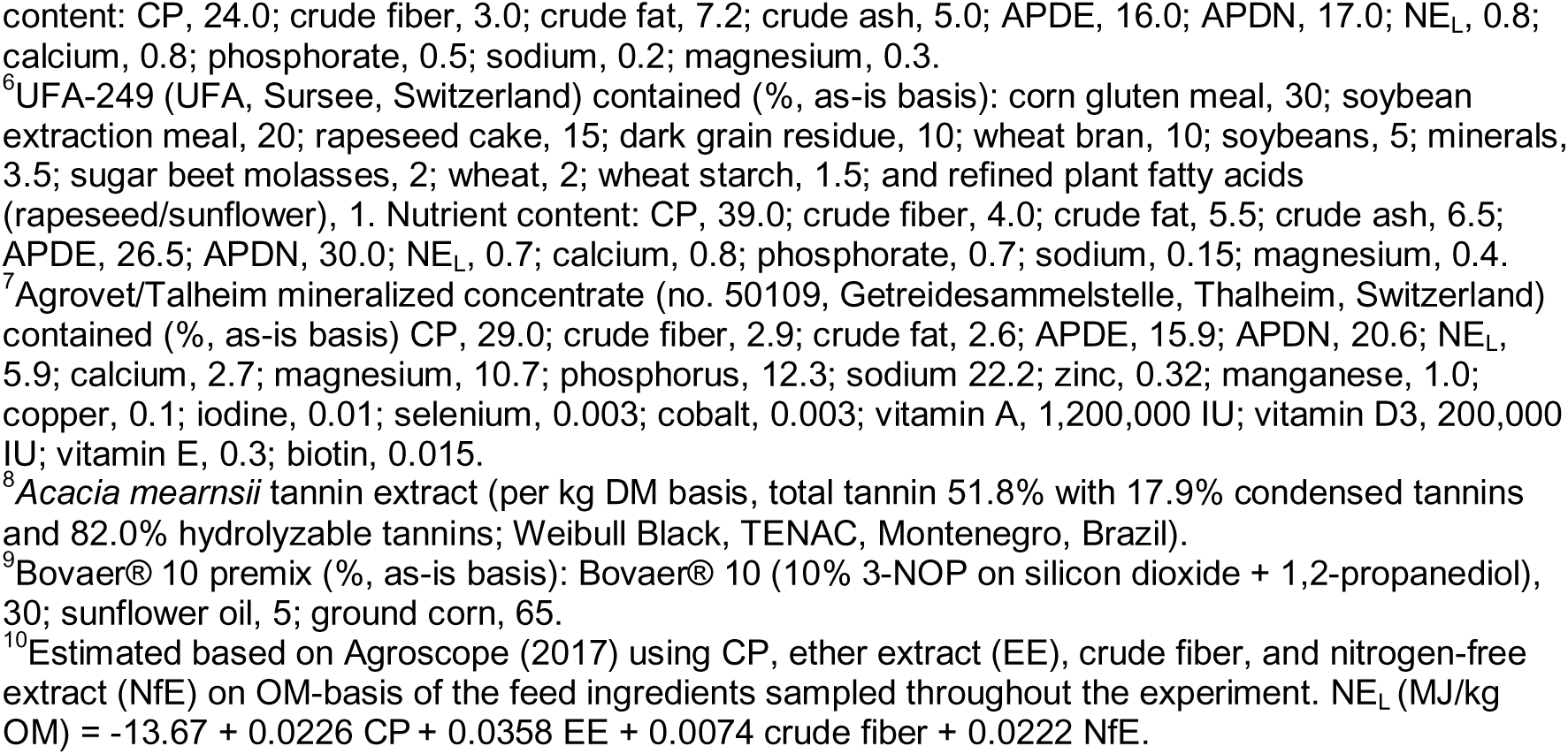
Ingredient and nutrient composition of the experimental diets.

| Items | Diets<sup>1</sup> |  |  |  |
| --- | --- | --- | --- | --- |
|  | CON | 3-NOP | TAN | 3-NOP + TAN |
| Feed ingredients, % of DM |  |  |  |  |
| Corn silage<sup>2</sup> | 37.9 | 37.4 | 37.2 | 32.9 |
| Grass silage<sup>3</sup> | 16.9 | 16.3 | 16.2 | 16.1 |
| Alfalfa hay<sup>4</sup> | 13.4 | 13.6 | 13.4 | 13.2 |
| Energy concentrate<sup>5</sup> | 10.9 | 10.5 | 10.4 | 13.7 |
| Protein concentrate<sup>6</sup> | 11.9 | 11.5 | 11.3 | 10.3 |
| Mineralized concentrate<sup>7</sup> | 7.02 | 7.01 | 6.96 | 7.40 |
| Ground corn | 1.60 | 1.59 | 1.54 | 1.70 |
| Tannin<sup>8</sup> | - | - | 3.00 | 3.00 |
| Bovaer® 10 premix<sup>9</sup> | - | 2.00 | - | 2.00 |
| Chemical composition, % of DM |  |  |  |  |
| CP | 15.9 | 15.8 | 15.9 | 15.9 |
| NDF | 40.6 | 40.4 | 40.7 | 40.5 |
| ADF | 21.6 | 20.9 | 21.6 | 21.6 |
| EE | 3.00 | 3.00 | 3.00 | 3.00 |
| Starch | 20.7 | 19.7 | 20.7 | 20.7 |
| Ash | 7.24 | 7.22 | 7.24 | 7.24 |
| NE<sub>L</sub>, MJ/kg of OM<sup>10</sup> | 5.40 | 5.30 | 5.40 | 5.40 |

<sup>1</sup>CON = basal diet; 3-NOP = 3-nitrooxypropanol supplemented diet (60 mg/kg DM); TAN = tannin extract (*Acacia mearnsii*) supplemented diet (3% of DM); 3-NOP + TAN = combination of 3-NOP and TAN supplemented diet.
<sup>2</sup>Corn silage was 35.5% DM and contained (DM basis) 6.30% CP and 52.5% NDF.
<sup>3</sup>Grass silage was 34.8% DM and contained (DM basis) 11.1 % CP and 65.1% NDF.
<sup>4</sup>Alfalfa hay was 90.5% DM and contained (DM basis) 12.3% CP and 53.0% NDF.
<sup>5</sup>UFA-173 (UFA, Sursee, Switzerland) contained (% as-is basis): soybean extraction meal, 25; corn flakes, 20; flaked barley (whole), 15; corn gluten meal, 8; wheat, 6; wheat flakes, 5; soybeans, 4; minerals, 4; crystalline fat (rapeseed), 3; sugar beet molasses, 3; rapeseed cake, 2; wheat starch, 2; dark grain residue, 1; refined plant fatty acids (rapeseed/sunflower), 1; and dextrose, 1. Nutrient

### Milk sampling and analyses

Cows were milked twice daily at 0530 and 1530 h. We recorded milk yields and collected samples during each milking from 20 to 24-d of each period using Bronopol-containing tubes. Samples were analyzed for fat, CP, lactose, and urea content following the International Organization for Standardization (ISO 9622) using a Fourier transform infrared spectrophotometer (MilkoScan RM 6 FT6000) at SuisseLab AG (Zollikofen, Switzerland). Milk urea nitrogen (**MUN**) concentration was calculated as urea concentration (mg/dL) × 0.466 (Beatson et al., 2019) and milk true protein as milk crude protein (%) − MUN (mg/dL) × 6.38 / 1000 (AOAC, 1997). Energy-corrected milk (**ECM**) yield was calculated based on the following equation (Sjaunja et al., 1990): ECM yield (kg/d) = 12.18 × Fat (kg/d) + 7.69 × Protein (kg/d) + 5.26 × Lactose (kg/d) + 0.01 × Milk Yield (kg/d).

### Feed sampling, analysis, and feeding behavior

Weekly grass and corn silage samples were collected and dried at 55°C for 48 h to adjust their DM content in the diet. Individual feed ingredients, TMR, orts, and GreenFeed pellets were sampled on d 18 and 20 of each experimental period and stored at –20°C. The samples were dried at 55°C for 48 h, and ground through a 1-mm sieve using a centrifugal mill (ZM 1, Retsch GmbH, Haan, Germany). For each period, TMR samples were composited by treatment and period, and orts by cow and period on an equal DM basis.

All samples were analyzed for DM, ash, NDF, ADF, N, starch, ether extract, and crude fiber according to the official methods of analysis (AOAC International, 1997) and performed in duplicate as described by Birkinshaw et al. (2022). Shortly, DM and total ash were determined using a thermogravimetric device (TGA-701, Leco; AOAC official method 942.5); organic matter (**OM**) was calculated as the difference between DM and total ash. Nitrogen was analyzed by a carbon/nitrogen analyzer (TruMac CN, Leco; AOAC official method 968.06). Crude protein was calculated as N × 6.25. Ether extract was determined using a Soxhlet extractor (Extraction System B-811, Büchi; AOAC official method 963.15), and a Fibertherm FT 12 (Gerhardt GmbH and Co. KG, Königswinter, Germany) to determine ash-corrected detergent fiber fractions. Heat-stable _α_-amylase (Sigma-Aldrich, St. Louis, MO) was used with NDF (AOAC official method 2002.04) and ADF (AOAC official method 973.18) analysis.

Feed weight was measured at 10-s intervals from d 18 to 24 of each period utilizing an automated feeding monitoring system (Mettler Toledo GmbH; Greifensee, Switzerland). The daily feed intake (**FI**) was quantified by subtracting the amount of refused feed from the feed offered. Measurements of hourly feed intake were determined by the continuous weight recordings of the feeders on an as-fed basis. The feed intake rate was expressed as the % of daily FI/h to standardize across cows with different total intake levels, allowing for relative comparisons of feeding behavior over time between BS and HF cows following a similar approach by Niu et al. (2014; 2017; 2018). Intra-day FI was calculated using an algorithm developed in the R statistical programming language (R Core Team 2022; version 4.3.3), according to Serviento et al. (2024).

### Fecal and urine sampling and analyses

Spot sampling of feces and urine was conducted from d 21 to 24 in each period for a total of 8 times at 1000, 1400 h, and 2200 h (sampling d 1), 0400, 1300, and 1800 h (sampling d 2), 0100 and 0700 h (sampling d 3). Fecal samples were collected either directly from the rectum or during spontaneous defecation. The samples were then sealed in plastic zip-lock bags and a set of fresh samples was combined for each cow and period to create a composite for total N analysis and kept at −20 °C until further processing. Similar to the feed samples, fecal samples were dried, ground, and composited based on the DM content for each cow and each period, and analyzed for DM, ash, NDF, ADF, N, starch, and acid-insoluble ash (**AIA**) contents as described above. The AIA content of the TMR and feces was used as an internal marker for total fecal DM output (Van Keulen and Young, 1977), to estimate the apparent total-tract digestibility (**ATTD**) of nutrients.

For urine, approximately 100 mL sample was collected by gently massaging the area below the vulva or capturing during spontaneous urination, and immediately after each collection, 10 mL of urine was mixed with 0.2 M sulfuric acid mix to reach a pH < 3 and composited by cow and period. The samples were stored at –20°C for further analysis. Composite urine samples were analyzed in-house for creatinine (DetectX® creatinine detection kit -K002-H1; Arbor Assays, Ann Arbor, MI), urinary urea nitrogen (**UUN**) (DetectX® urea nitrogen kit - K024-H1; Arbor Assays, Ann Arbor, MI), and uric acid (Abbexa®; Catalog: abx298824, Science Park, Cambridge, UK) using a chromate microplate reader (Cytation3, BioTek’s Hybrid Technology, USA) set at a wavelength of 490 and 450 nm, respectively. Allantoin was analyzed according to Chen et al. (1992) and read at wavelengths of 522 nm with a UV/visible spectrophotometer (Beckman Coulter Inc., California, USA). Urine volume was estimated from the creatinine concentration in urine assuming a constant daily creatinine excretion rate of 29 mg/dL per kg of BW (Hristov et al., 2010). Excretion of allantoin, uric acid, total purine derivatives (**PD**, allantoin + uric acid), and total N in urine was calculated by multiplying the concentration of each metabolite by the estimated daily urine volume.

### Enteric gas emission

Enteric emissions of CH_4_, hydrogen (**H_2_**), and carbon dioxide (**CO_2_**) were measured from individual animals using a single unit of GreenFeed System (GF; C-Lock, Inc., Rapid City, SD, USA) 8 times during the last 3 d of each experimental period to represent a 24-h feeding cycle: 0900, 1500, and 2100 h (sampling d 1), 0300, 1200, and 1900 h (sampling d 2), 0000 and 0600 h (sampling d 3) based on method adopted from Hristov et al. (2015). Cows were trained to use GF 10-d before the start of the experiment while in the free-stall barn. The GF feeding schedule was configured to allow a maximum of 8 bait drops (approximately 300 g/event per cow) of dehydrated alfalfa pellets (Luzatop, Désialis, Paris, France) with an 18% CP content (on a DM-basis) for a continuous measurement of 5 min/cow. Pellet consumption was included in the daily DMI calculation during sampling days. Each cow was assigned a collar-mounted radio-frequency identification tag for reading by the GF system. Before each measurement, the GF unit underwent calibration using a standard zero and span baseline gas (Linde Gas Benelux B.V.). To ensure accurate background air measurement, the GF was moved away from the cow for a 2-min washout between measurements. The total measurement duration, including washout, of the 16 cows at each sampling was less than 2 h. A CO_2_ recovery test was conducted before each sampling session. Data processing was managed by C-Lock Inc., and statistical analyses were conducted on the validated data set.

### Statistical analysis

Data analysis and calculations, including DMI, milk yield, ATTD of nutrients, N partitioning, feeding, and gas emissions, were conducted in R using the mixed-model approach with the *lmer* procedure (Bates et al., 2015). The Kenward-Roger method was applied to adjust the denominator degrees of freedom for more accurate statistical inference. The full model was as follows:

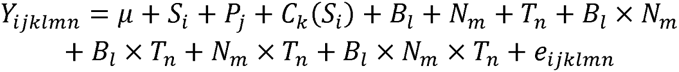

where *Y_i jklmn_* is variable of interest, *μ* is the overall mean, *S_i_* is random effect of sequence (*i* = 1–4), *P_j_* is the fixed effect of period (*j* = 1–4), *C_k_*(*S_i_*) is the random effect of cow nested in sequence (*k* = 1–16), *B_l_* is the fixed effect of main plot (breed of cows; *l* = 1–2), *N_m_* is the fixed effect of 3-NOP (*m* = 1–2), *T_n_* is the fixed effect of TAN (*n* = 1–2), *B_l_* ×*N_m_* is the interaction of breed by 3-NOP, *B_l_* ×*T_n_* is the interaction of the breed by TAN, *N_m_* ×*T_n_* is the interaction of 3-NOP by TAN, *B_l_* × *N_m_* ×*T_n_*. is the 3-way interaction of breed, 3-NOP, and TAN, and *e_i jklmn_* is the random residual.

For enteric CH_4_ production, there was no 3-way interaction; however, breed by 3-NOP interaction was shown. Therefore, the enteric gas emission data at 3-h intervals and hourly feeding behavior were analyzed using a mixed model with repeated measures of time, employing the *lmer* procedure (Bates et al., 2015). The model was as follows:

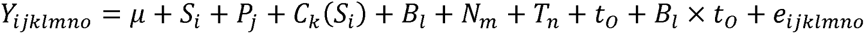

where *Y_i jklmno_* is the intra-day gas emission or feeding behavior data,*t_o_* is the time of day (*o* = 1–8 for enteric gas emission and 1–24 for feeding behavior data, respectively). Other terms in the model were the same as described above.

Data points exhibiting studentized residuals beyond ±3 were considered outliers and excluded from the analysis, with no more than 2 entries removed from each parameter. Specifically, 8 observations from all variables in 8 cows (4 from each breed HF and BS) were omitted from the first period due to a calculation error with dosing of TAN, and during the last period, one cow was no longer part of the experiment due to health issues unrelated to the experiment. Pre-planned contrasts tested were 3-NOP by breed, and 3-NOP by TAN interactions. Treatment means were compared using the Tukey-Kramer method, with significance defined at *P* ≤ 0.05 for main effects and *P* ≤ 0.10 for interactions. Tendencies were identified at 0.05 < *P* ≤ 0.10 for main effects and 0.10 < *P* ≤ 0.15 for interactions (Piantoni et al., 2015; de Souza et al., 2019). Non-significant interaction terms were removed, leading to data being adjusted and presented as least-squares means (**LSM**) from the simplified model without these terms.

## Results

Three-way interactions among 3-NOP, TAN, and Breed were non-significant for all responses except for CH_4_ yield and CO_2_ production. Therefore, the following results will emphasize two-way interactions and the main effects of individual treatments.

### Milk production

Data on lactational performance is summarized in Table 2, and treatment level data is provided in Supplementary Table S1. As expected, HF cows, on average, consumed approximately 3.7 kg/d more (*P* < 0.001) DMI than the BS cows, and similarly, HF cows produced around 9.3 kg/d more (*P* < 0.001) milk than BS, resulting in greater feed efficiency for HF, as expressed by ECM:DMI ratio (*P* = 0.016). Dry matter intake was affected by TAN (*P* = 0.003) with a reduction of 1 kg/d. A tendency was observed for interaction between 3-NOP and TAN for DMI, as cows fed TAN alone tended to have (*P =* 0.14) a greater reduction in DMI. Neither 3-NOP nor TAN affected milk yield or ECM. However, statistical tendencies were observed for 3-NOP × Breed, where 3-NOP increased milk yield (*P =* 0.13) and ECM (*P =* 0.12) in HF cows but decreased them in BS cows. Further, a tendency for a 3-NOP × TAN interaction in fat yield (*P* = 0.11) and fat concentration (*P* = 0.15) was observed where fat yield tended to be greater with 3-NOP supplementation. Feeding TAN tended (*P* = 0.062) to reduce milk lactose yield, and 3-NOP tended (*P* = 0.088) to increase milk lactose concentration. A 3-NOP × Breed interaction (*P* = 0.083) was observed for lactose yield because only HF cows supplemented with 3-NOP increased it. Cows fed TAN had 18.6% lower (*P* < 0.001) MUN concentration. Milk true protein yield tended (*P =* 0.061) to reduce, whereas its concentration decreased (*P =* 0.054) in cows fed TAN.

**Table 2.**
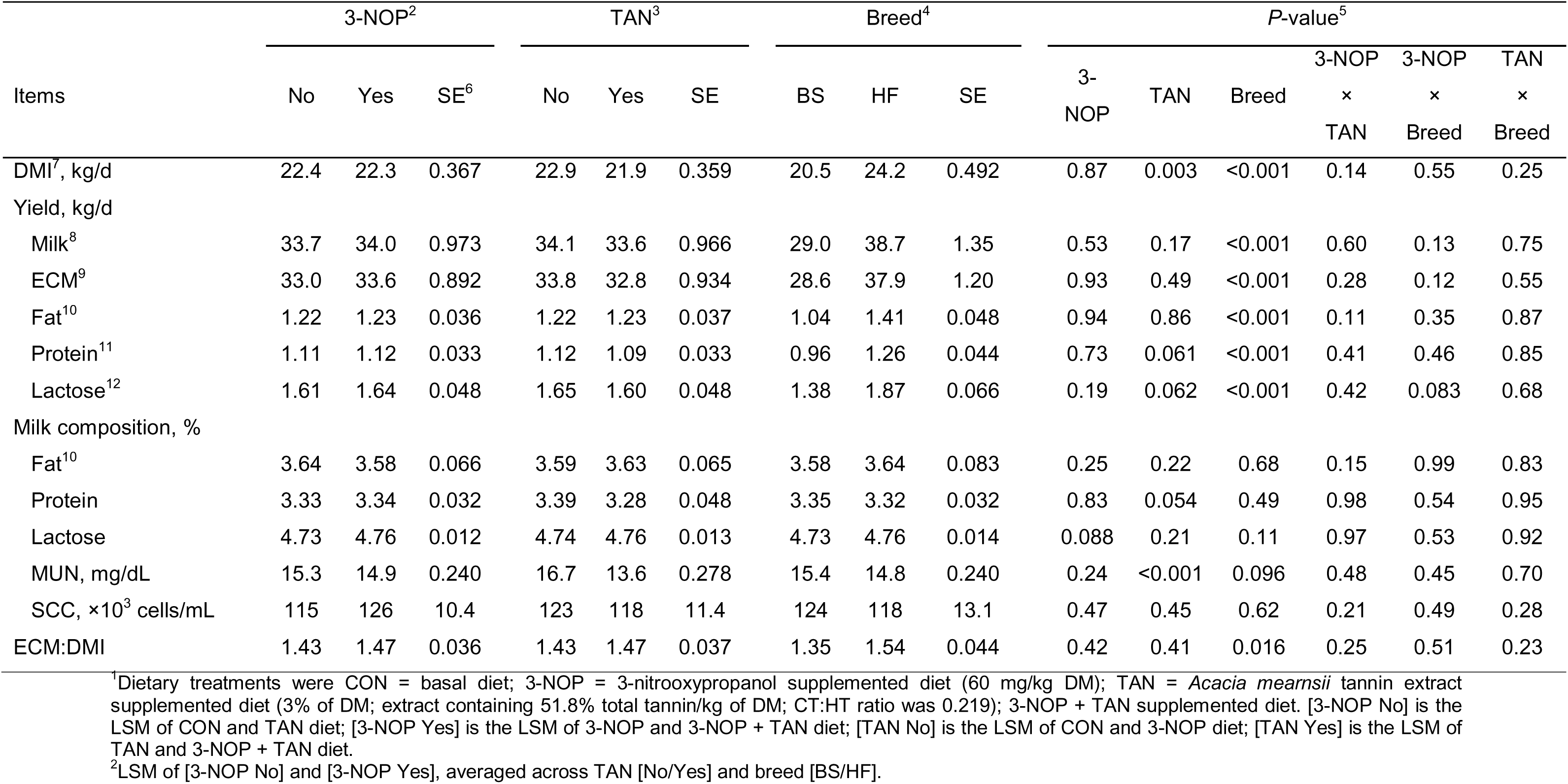
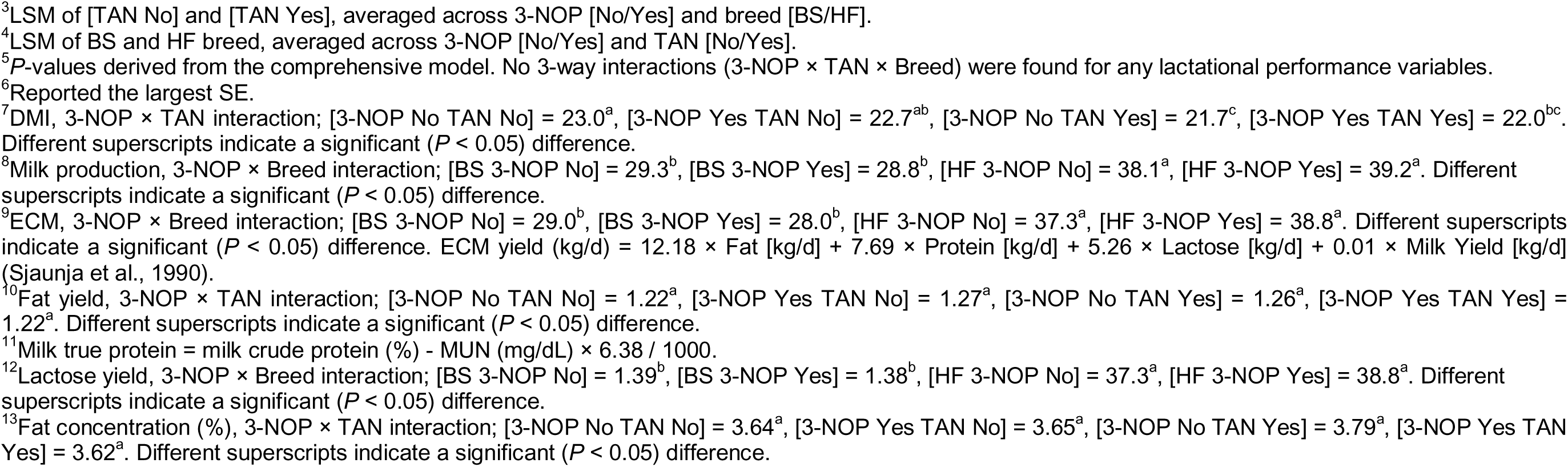
Lactational performance of Brown Swiss (BS) and Holstein Friesian (HF) cows fed dietary treatments^1^.

| Items | 3-NOP<sup>2</sup> |  |  | TAN<sup>3</sup> |  |  | Breed<sup>4</sup> |  |  | P-value<sup>5</sup> |  |  |  |  |  |
| --- | --- | --- | --- | --- | --- | --- | --- | --- | --- | --- | --- | --- | --- | --- | --- |
|  | No | Yes | SE<sup>6</sup> | No | Yes | SE | BS | HF | SE | 3-NOP | TAN | Breed | 3-NOP | 3-NOP | TAN |
|  |  |  |  |  |  |  |  |  |  |  |  |  | ×<br>TAN | ×<br>Breed | ×<br>Breed |
| DMI<sup>7</sup>, kg/d | 22.4 | 22.3 | 0.367 | 22.9 | 21.9 | 0.359 | 20.5 | 24.2 | 0.492 | 0.87 | 0.003 | <0.001 | 0.14 | 0.55 | 0.25 |
| Yield, kg/d |  |  |  |  |  |  |  |  |  |  |  |  |  |  |  |
| Milk<sup>8</sup> | 33.7 | 34.0 | 0.973 | 34.1 | 33.6 | 0.966 | 29.0 | 38.7 | 1.35 | 0.53 | 0.17 | <0.001 | 0.60 | 0.13 | 0.75 |
| ECM<sup>9</sup> | 33.0 | 33.6 | 0.892 | 33.8 | 32.8 | 0.934 | 28.6 | 37.9 | 1.20 | 0.93 | 0.49 | <0.001 | 0.28 | 0.12 | 0.55 |
| Fat<sup>10</sup> | 1.22 | 1.23 | 0.036 | 1.22 | 1.23 | 0.037 | 1.04 | 1.41 | 0.048 | 0.94 | 0.86 | <0.001 | 0.11 | 0.35 | 0.87 |
| Protein<sup>11</sup> | 1.11 | 1.12 | 0.033 | 1.12 | 1.09 | 0.033 | 0.96 | 1.26 | 0.044 | 0.73 | 0.061 | <0.001 | 0.41 | 0.46 | 0.85 |
| Lactose<sup>12</sup> | 1.61 | 1.64 | 0.048 | 1.65 | 1.60 | 0.048 | 1.38 | 1.87 | 0.066 | 0.19 | 0.062 | <0.001 | 0.42 | 0.083 | 0.68 |
| Milk composition, % |  |  |  |  |  |  |  |  |  |  |  |  |  |  |  |
| Fat<sup>10</sup> | 3.64 | 3.58 | 0.066 | 3.59 | 3.63 | 0.065 | 3.58 | 3.64 | 0.083 | 0.25 | 0.22 | 0.68 | 0.15 | 0.99 | 0.83 |
| Protein | 3.33 | 3.34 | 0.032 | 3.39 | 3.28 | 0.048 | 3.35 | 3.32 | 0.032 | 0.83 | 0.054 | 0.49 | 0.98 | 0.54 | 0.95 |
| Lactose | 4.73 | 4.76 | 0.012 | 4.74 | 4.76 | 0.013 | 4.73 | 4.76 | 0.014 | 0.088 | 0.21 | 0.11 | 0.97 | 0.53 | 0.92 |
| MUN, mg/dL | 15.3 | 14.9 | 0.240 | 16.7 | 13.6 | 0.278 | 15.4 | 14.8 | 0.240 | 0.24 | <0.001 | 0.096 | 0.48 | 0.45 | 0.70 |
| SCC, ×10<sup>3</sup> cells/mL | 115 | 126 | 10.4 | 123 | 118 | 11.4 | 124 | 118 | 13.1 | 0.47 | 0.45 | 0.62 | 0.21 | 0.49 | 0.28 |
| ECM:DMI | 1.43 | 1.47 | 0.036 | 1.43 | 1.47 | 0.037 | 1.35 | 1.54 | 0.044 | 0.42 | 0.41 | 0.016 | 0.25 | 0.51 | 0.23 |

<sup>1</sup>Dietary treatments were CON = basal diet; 3-NOP = 3-nitrooxypropanol supplemented diet (60 mg/kg DM); TAN = *Acacia mearnsii* tannin extract supplemented diet (3% of DM; extract containing 51.8% total tannin/kg of DM; CT:HT ratio was 0.219); 3-NOP + TAN supplemented diet. [3-NOP No] is the LSM of CON and TAN diet; [3-NOP Yes] is the LSM of 3-NOP and 3-NOP + TAN diet; [TAN No] is the LSM of CON and 3-NOP diet; [TAN Yes] is the LSM of TAN and 3-NOP + TAN diet.
<sup>2</sup>LSM of [3-NOP No] and [3-NOP Yes], averaged across TAN [No/Yes] and breed [BS/HF].

### Apparent total-tract digestibility, N utilization, and purine derivatives

Data regarding ATTD are outlined in Table 3, and treatment level data is provided in Supplementary Table S2. The ATTD of OM and ADF did not differ between cow breeds fed 3-NOP or TAN. An interaction between TAN and Breed tended to affect NDF digestibility (*P* = 0.067), with TAN reducing NDF digestibility by 4.47% in BS and by 26.5% in HF cows. Additionally, a tendency for 3-NOP × Breed interaction was observed (*P* = 0.12), where 3-NOP increased 23.6% NDF digestibility in BS but decreased 5.38% in HF cows. The ATTD of CP tended to be lower (*P* = 0.080) for cows fed TAN, and differed between breeds, with HF showing a greater ATTD of CP (*P* = 0.050) compared to BS.

**Table 3.**
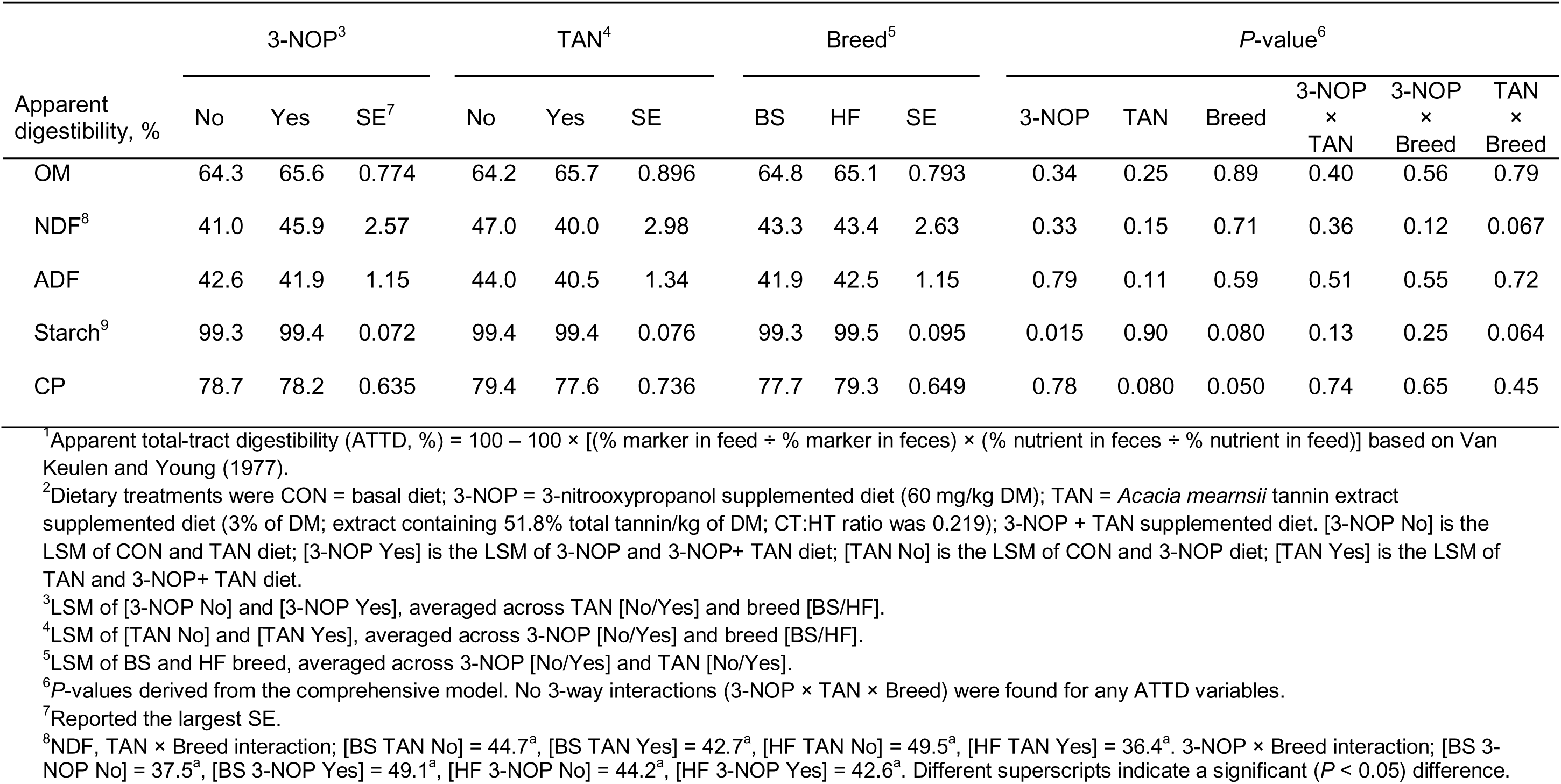
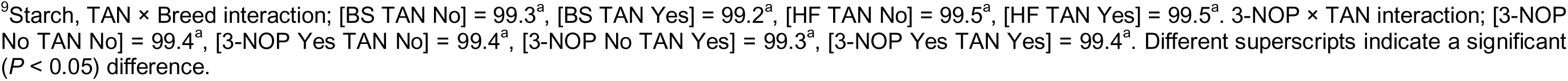
Apparent total-tract digestibility (ATTD)^1^ of Brown Swiss (BS) and Holstein Frisian (HF) cows fed dietary treatments^2^.

| Apparent digestibility, % | 3-NOP<sup>3</sup> |  |  | TAN<sup>4</sup> |  |  | Breed<sup>5</sup> |  |  | P-value<sup>6</sup> |  |  |  |  |  |
| --- | --- | --- | --- | --- | --- | --- | --- | --- | --- | --- | --- | --- | --- | --- | --- |
|  | No | Yes | SE<sup>7</sup> | No | Yes | SE | BS | HF | SE | 3-NOP | TAN | Breed | 3-NOP<br>×<br>TAN | 3-NOP<br>×<br>Breed | TAN<br>×<br>Breed |
| OM | 64.3 | 65.6 | 0.774 | 64.2 | 65.7 | 0.896 | 64.8 | 65.1 | 0.793 | 0.34 | 0.25 | 0.89 | 0.40 | 0.56 | 0.79 |
| NDF<sup>8</sup> | 41.0 | 45.9 | 2.57 | 47.0 | 40.0 | 2.98 | 43.3 | 43.4 | 2.63 | 0.33 | 0.15 | 0.71 | 0.36 | 0.12 | 0.067 |
| ADF | 42.6 | 41.9 | 1.15 | 44.0 | 40.5 | 1.34 | 41.9 | 42.5 | 1.15 | 0.79 | 0.11 | 0.59 | 0.51 | 0.55 | 0.72 |
| Starch<sup>9</sup> | 99.3 | 99.4 | 0.072 | 99.4 | 99.4 | 0.076 | 99.3 | 99.5 | 0.095 | 0.015 | 0.90 | 0.080 | 0.13 | 0.25 | 0.064 |
| CP | 78.7 | 78.2 | 0.635 | 79.4 | 77.6 | 0.736 | 77.7 | 79.3 | 0.649 | 0.78 | 0.080 | 0.050 | 0.74 | 0.65 | 0.45 |

<sup>1</sup>Apparent total-tract digestibility (ATTD, %) = 100 – 100 × [(% marker in feed ÷ % marker in feces) × (% nutrient in feces ÷ % nutrient in feed)] based on Van Keulen and Young (1977).
<sup>2</sup>Dietary treatments were CON = basal diet; 3-NOP = 3-nitrooxypropanol supplemented diet (60 mg/kg DM); TAN = *Acacia mearnsii* tannin extract supplemented diet (3% of DM; extract containing 51.8% total tannin/kg of DM; CT:HT ratio was 0.219); 3-NOP + TAN supplemented diet. [3-NOP No] is the LSM of CON and TAN diet; [3-NOP Yes] is the LSM of 3-NOP and 3-NOP+ TAN diet; [TAN No] is the LSM of CON and 3-NOP diet; [TAN Yes] is the LSM of TAN and 3-NOP+ TAN diet.
<sup>3</sup>LSM of [3-NOP No] and [3-NOP Yes], averaged across TAN [No/Yes] and breed [BS/HF].
<sup>4</sup>LSM of [TAN No] and [TAN Yes], averaged across 3-NOP [No/Yes] and breed [BS/HF].
<sup>5</sup>LSM of BS and HF breed, averaged across 3-NOP [No/Yes] and TAN [No/Yes].
<sup>6</sup>P-values derived from the comprehensive model. No 3-way interactions (3-NOP × TAN × Breed) were found for any ATTD variables.
<sup>7</sup>Reported the largest SE.
<sup>8</sup>NDF, TAN × Breed interaction; [BS TAN No] = 44.7<sup>a</sup>, [BS TAN Yes] = 42.7<sup>a</sup>, [HF TAN No] = 49.5<sup>a</sup>, [HF TAN Yes] = 36.4<sup>a</sup>. 3-NOP × Breed interaction; [BS 3- NOP No] = 37.5<sup>a</sup>, [BS 3-NOP Yes] = 49.1<sup>a</sup>, [HF 3-NOP No] = 44.2<sup>a</sup>, [HF 3-NOP Yes] = 42.6<sup>a</sup>. Different superscripts indicate a significant (*P* < 0.05) difference.
<sup>9</sup>Starch, TAN × Breed interaction; [BS TAN No] = 99.3<sup>a</sup>, [BS TAN Yes] = 99.2<sup>a</sup>, [HF TAN No] = 99.5<sup>a</sup>, [HF TAN Yes] = 99.5<sup>a</sup>. 3-NOP × TAN interaction; [3-NOP No TAN No] = 99.4<sup>a</sup>, [3-NOP Yes TAN No] = 99.4<sup>a</sup>, [3-NOP No TAN Yes] = 99.3<sup>a</sup>, [3-NOP Yes TAN Yes] = 99.4<sup>a</sup>. Different superscripts indicate a significant ( $P < 0.05$ ) difference.

Data regarding N partitioning and PD excretions are presented in Table 4, and treatment level data is provided in Supplementary Table S3. There was no effect of 3-NOP supplementation on N utilization or PD excretions. Cows fed TAN had a tendency (*P =* 0.077) for reduced N intake. An interaction of TAN × Breed was observed (*P =* 0.013), indicating that TAN reduced urinary N excretion by 25.8% in BS and 20.2% in HF cows. Further, TAN supplementation resulted in a shift in N excretion from urine to feces, with urinary N excretion reduced by 23.0% (*P* < 0.001) and fecal N excretion increased by 36.7% (*P* < 0.001) across cow breeds. A tendency for 3-NOP × TAN interaction (*P* = 0.14) was observed, as cows fed only TAN had a greater reduction in urinary N (g/d) excretion. In addition, TAN reduced UUN excretion (*P* < 0.001) and decreased (*P* = 0.006) milk N (g/d). A 3-NOP × Breed interaction affected uric acid excretion (*P* = 0.014), with 3-NOP decreasing excretion by 11.9 mmol/d in BS but increasing by 13.7 mmol/d in HF cows. Neither 3-NOP nor TAN affected allantoin or total PD excretions.

**Table 4.**
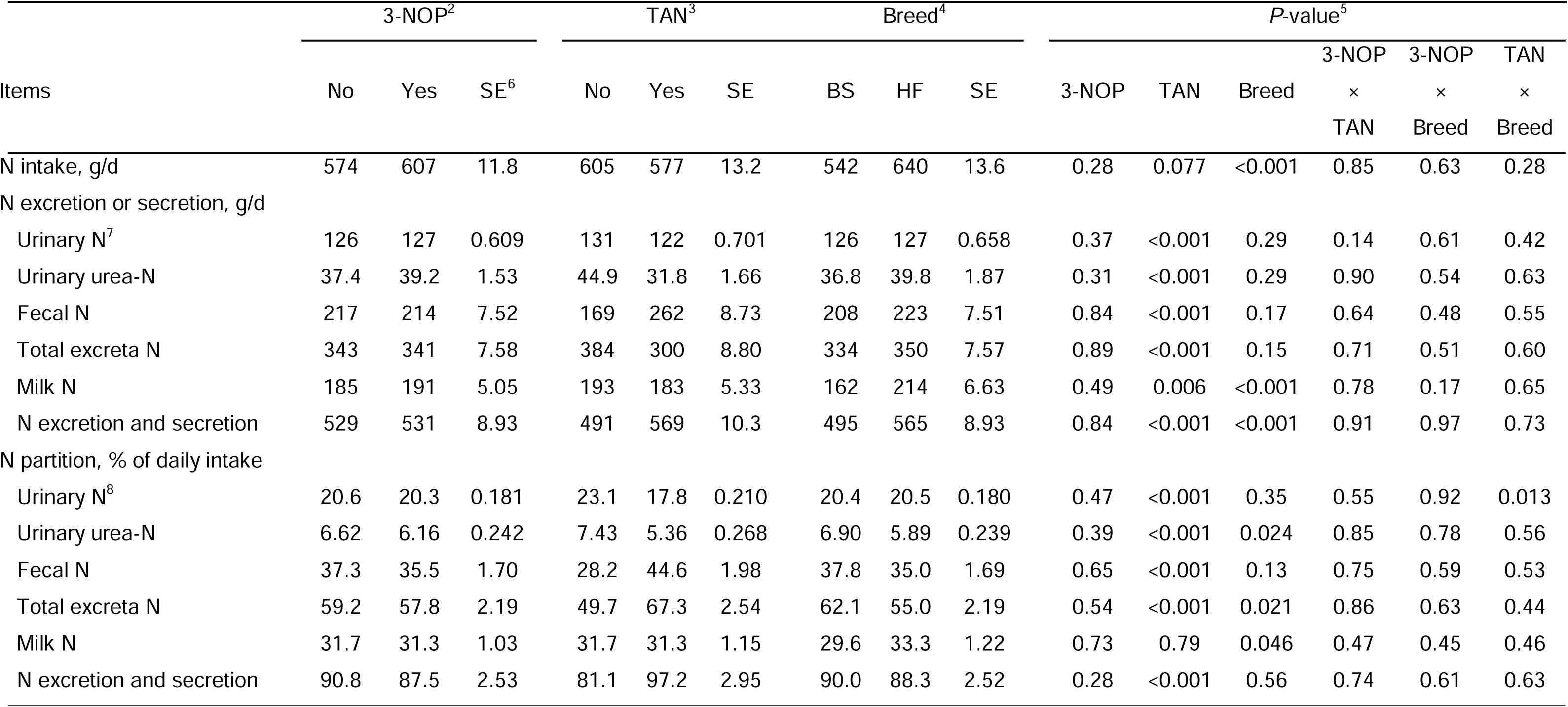
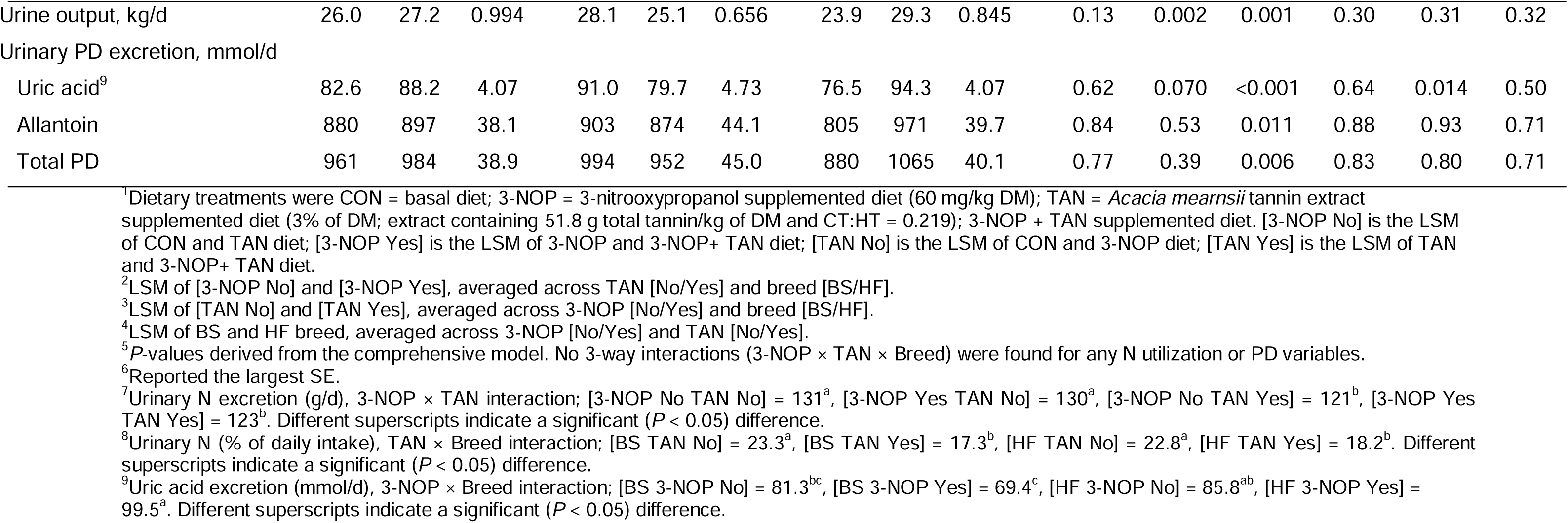
Nitrogen utilization and purine derivatives (PD) excretion in Brown Swiss (BS) and Holstein Friesian (HF) cows fed dietary treatments^1^.

### Enteric gas emissions

Enteric CH_4_, H_2_, and CO_2_ emissions are depicted in Table 5 and Figure 1, and treatment level data is provided in Supplementary Table S4. No 3-NOP × TAN interaction was observed for any gas variables except for CO_2_ production (*P* = 0.032). The reduction in CH_4_ production was influenced by an interaction (*P* = 0.050) between 3-NOP and breed, with BS cows showing a 13.0% reduction and HF cows exhibiting a more pronounced 21.6% reduction when supplemented with 3-NOP (Figure 1A). A 3-NOP × TAN × Breed interaction was observed for CH_4_ yield (*P* = 0.10) where HF cows decreased CH_4_ yield by 24.9% with 3-NOP alone, whereas in BS cows, CH_4_ yield was greatest with TAN alone and decreased by 4.74% with 3-NOP, the combination of 3-NOP and TAN did not enhance CH_4_ mitigation efficacy. A tendency of 3-NOP × Breed interaction was observed in enteric CH_4_ yield (g/kg DMI) and intensity (g/kg ECM) because HF cows had greater reductions (21.8% and 23.4%, respectively; Figure 1C) efficacy when fed 3-NOP compared to BS cows (11.0% and 10.8%, respectively; Figure 1D). Moreover, a tendency for TAN × Breed interaction was observed for CH_4_ intensity (g/kg milk yield; *P* = 0.12), because TAN increased CH_4_ intensity by 4.5% in BS, whereas it decreased by 5.1% in HF cows.

**Fig. 1.**
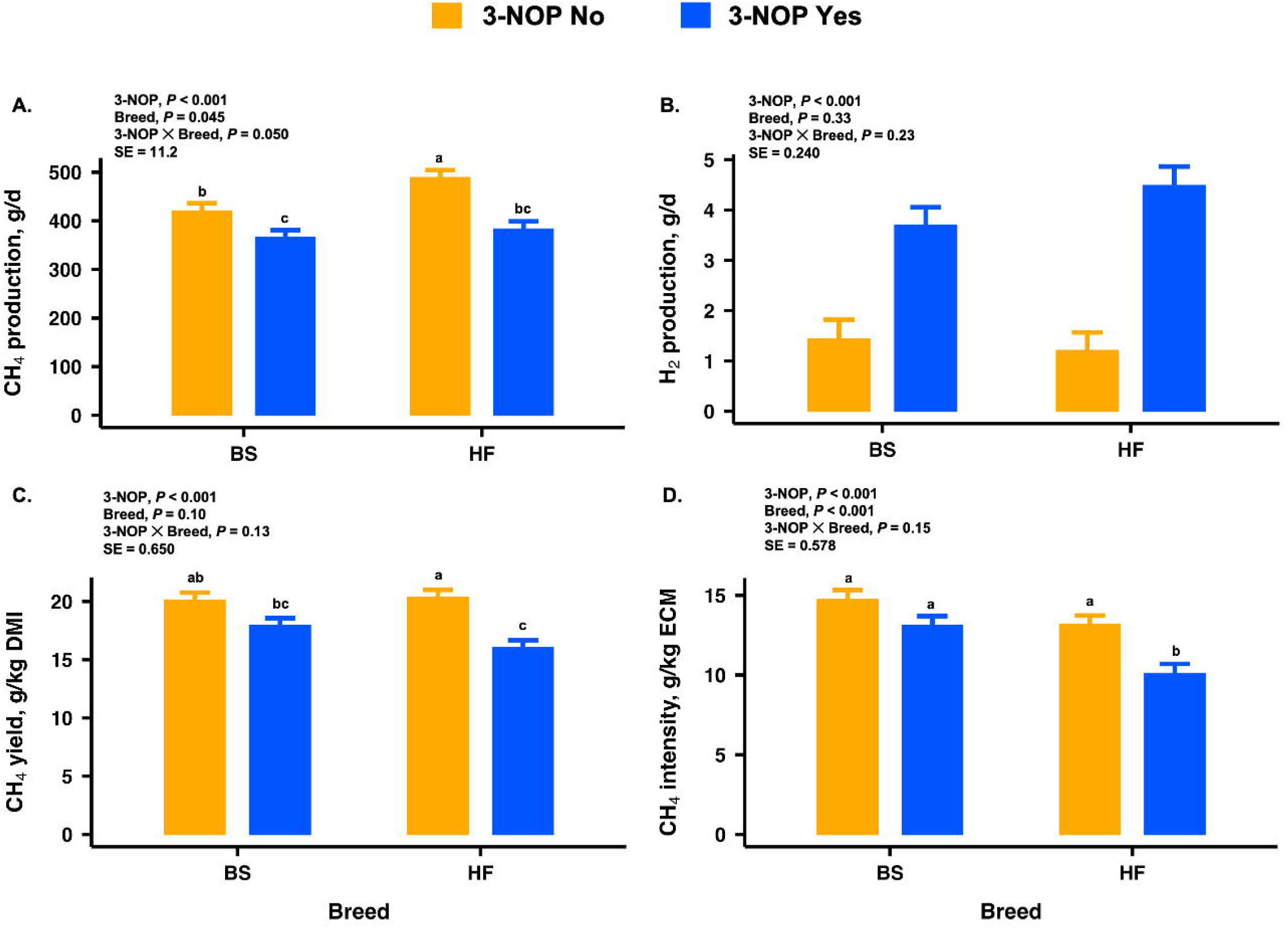
Interactions of 3-NOP and dairy cattle breeds [Brown Swiss (BS) and Holstein Friesian (HF)] on the CH_4_ production (A), H_2_ production (B), CH_4_ yield (C), and CH_4_ intensity (D). Dietary treatments were CON = basal diet; 3-NOP supplemented diet at 60 mg/kg DM; *Acacia mearnsii* tannin extract (TAN, 3% of DM); 3-NOP + TAN supplemented diet. [3-NOP No] is the LSM of CON and TAN diet; [3-NOP Yes] is the LSM of 3-NOP and 3-NOP + TAN diet. Data are presented as LSM ± SE.

**Table 5.**
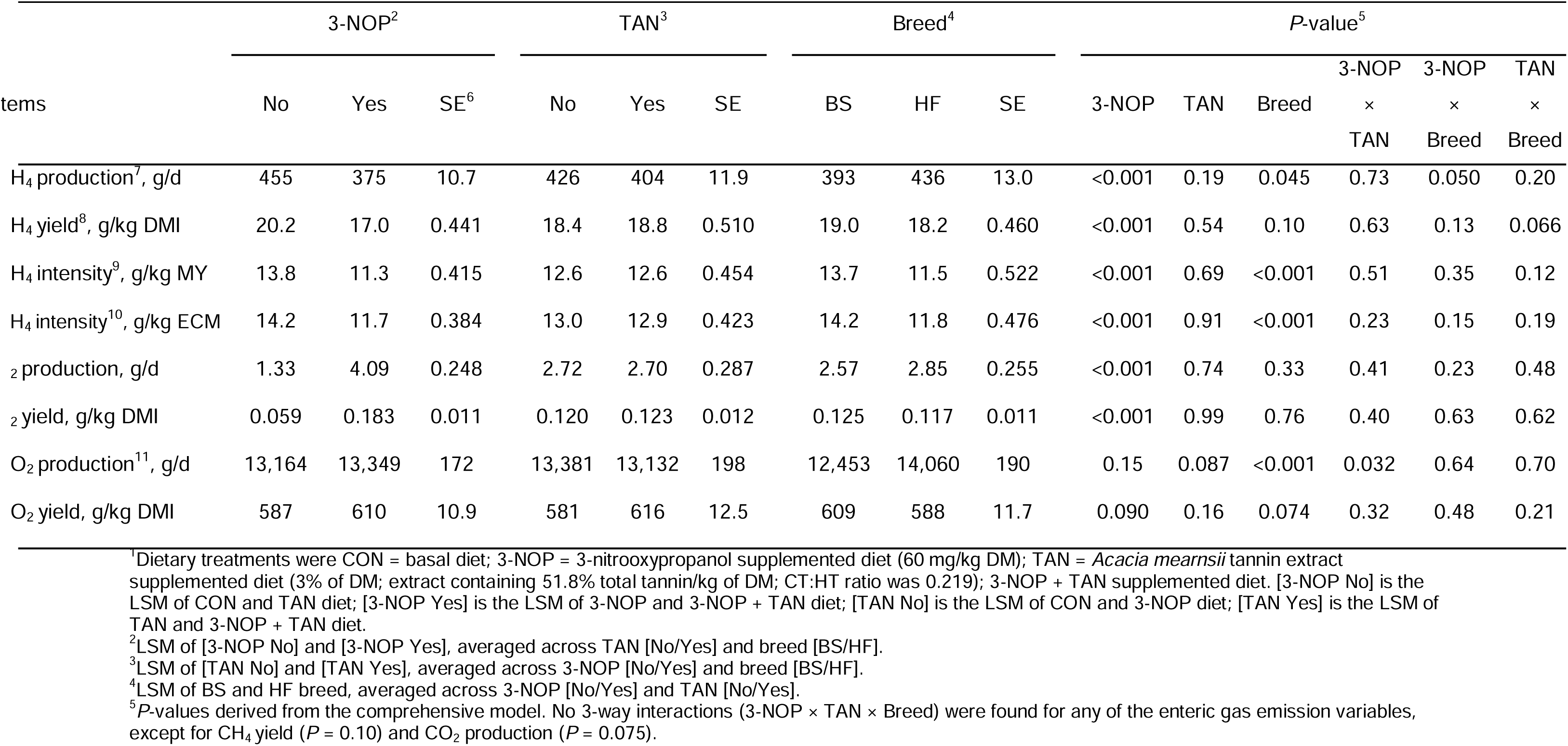
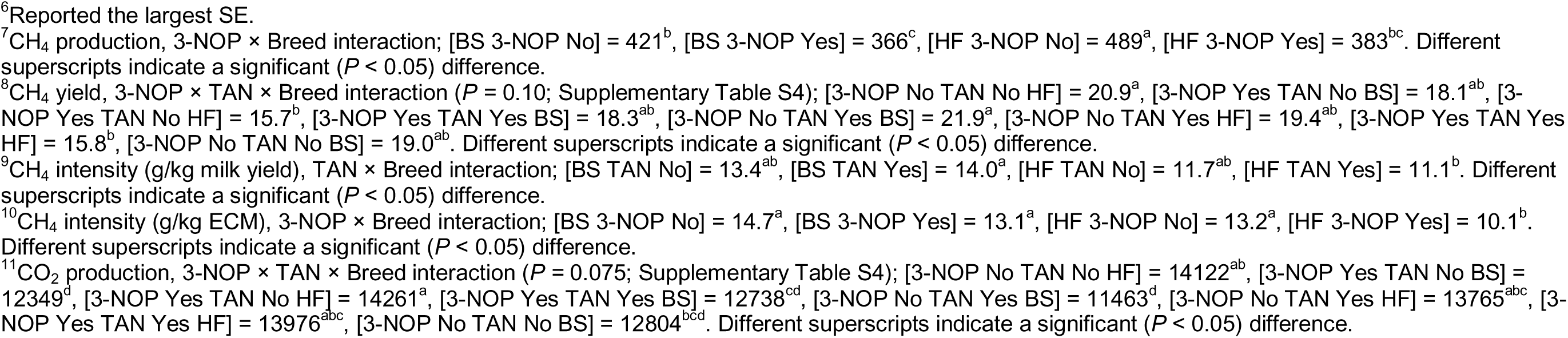
Enteric gas emissions of Brown Swiss (BS) and Holstein Friesian (HF) cows fed dietary treatments^1^.

Contrary to CH_4_ production, H_2_ showed no 3-NOP × Breed interaction (Figure 1B). Hydrogen production was increased (*P* < 0.001) with 3-NOP supplementation, while TAN supplementation had no effect on H_2_ emissions. There was a 3-NOP × TAN × Breed interaction (*P* = 0.075) for CO_2_ production, indicating the lowest CO_2_ production in BS cows with TAN, whereas HF cows supplemented with 3-NOP tended to greater CO_2_ production. The CO_2_ yield tended to increase (*P* = 0.090) in cows with 3-NOP supplementation.

### Diurnal pattern of feed intake, enteric CH_4,_ and H_2_ production

The diurnal variations in feed intake, CH_4_, and H_2_ production across a 24-h feeding cycle are illustrated in Figure 2, with LSM presented in Supplementary Tables S5-S7. Neither 3-NOP nor its interaction with time affected the daily FI patterns in either breed (Figure 2E and 2F). However, daily FI fluctuated across the day (*P* < 0.001), with BS and HF cows consuming 10.9% and 12.3%, respectively, within 1-h post-morning feeding, and 11.0% post-evening feeding. The HF cows had greater morning FI, however, no notable FI difference was observed between morning and evening feedings for both breeds.

**Fig. 2.**
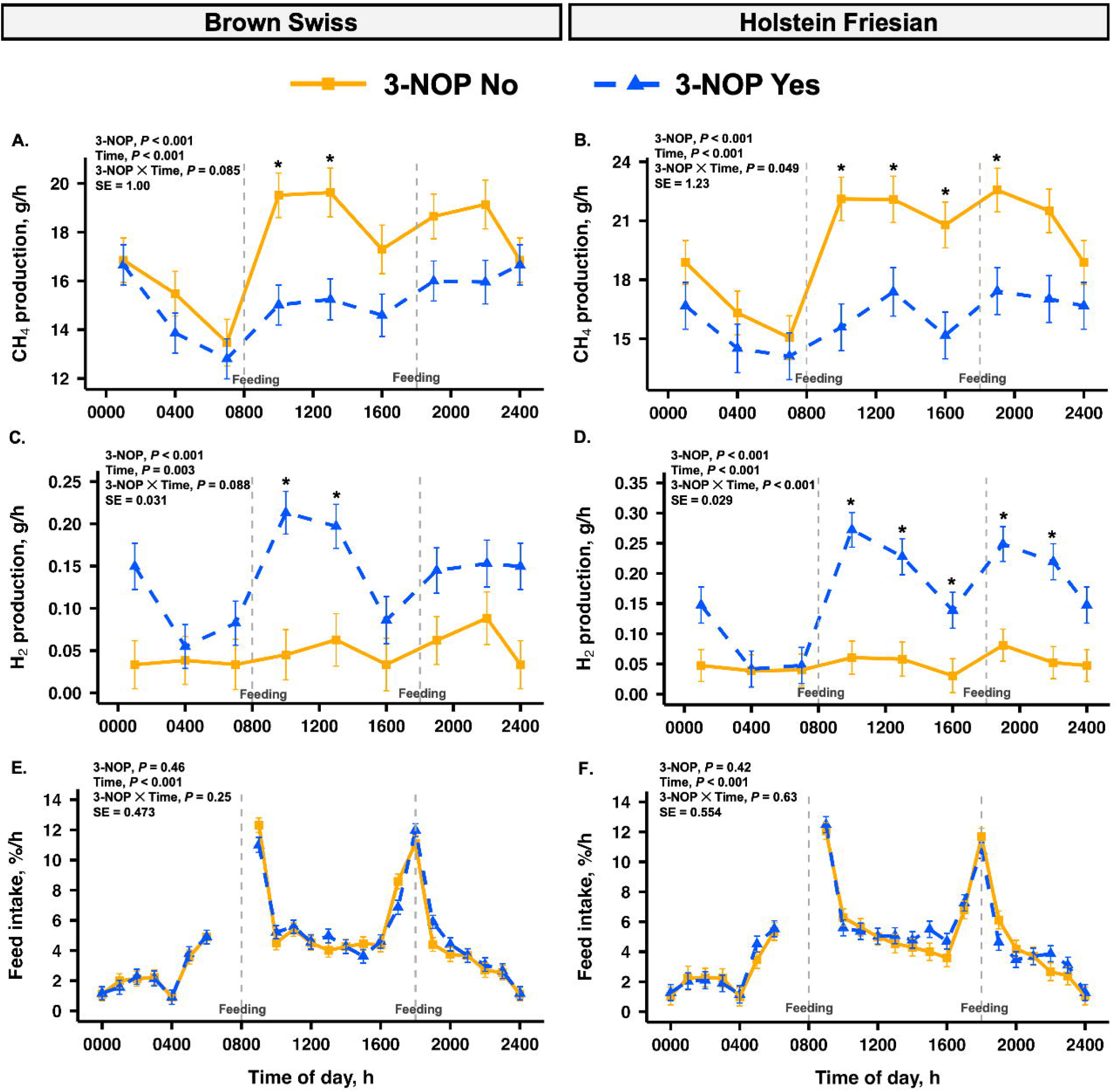
Diurnal variation in CH_4_ (A and B) and H_2_ (C and D) production (3-h intervals), and feed intake (E and F; hourly) for Brown Swiss (left panel) and Holstein Friesian (right panel) cows. Dietary treatments were CON = basal diet; 3-NOP supplemented diet at 60 mg/kg DM; *Acacia mearnsii* tannin extract (TAN, 3% of DM); 3-NOP + TAN supplemented diet. [3-NOP No] is the LSM of CON and TAN diet; [3-NOP Yes] is the LSM of 3-NOP and 3-NOP + TAN diet. Grey dashed lines mark feeding times at 0800 and 1800 h. No 3-way interaction of 3-NOP × Breed × Time was observed.

Additionally, there was a 3-NOP × Time interaction for CH_4_ production for HF (*P* = 0.049) and for BS cows (*P* = 0.085) (Figure 2A and 2B). The HF and BS cows without 3-NOP supplementation had increased CH_4_ production following morning and afternoon feeding. In contrast, the increase in CH_4_ production was less pronounced in cows supplemented with 3-NOP. The greatest intraday absolute CH_4_ reductions in BS cows were observed after the morning feeding from 1000 to 1300 h, averaging 22.8%, whereas HF cows exhibited a higher and more consistent reduction from 1000 to 1900 h, averaging 25.2% (Figure 2A and 2B; Supplementary Table S5). The mitigating effects of 3-NOP during the late night and early morning periods were relatively low, regardless of animal breed. Overall, the mitigating effect was most pronounced immediately after morning feeding, continued strongly until midnight, diminished overnight, and was at its lowest before the next day’s morning feeding.

Similarly, an interaction between 3-NOP and time was observed for H_2_ production for HF (*P* < 0.001), and BS (*P* = 0.088) cows, with 3-NOP supplemented cows having a greater H_2_ production from 1000 h to 2200 h. The most notable increases in H_2_ production, 371% for BS and 355% for HF occurred at 1000 h (*P* < 0.05; Figure 2C and 2D; Supplementary Table S6), while no noticeable differences were detected between treatment groups before the morning feed around 0700 h. When 3-NOP was fed, H_2_ production of BS cows increased following morning feeding (1000 and 1300 h, on average by 2.9 folds), whereas both morning (1000 and 1300 h, on average by 3.3 folds) and afternoon (1900 and 2200 h, on average by 2.7 folds) feeding triggered increment in HF cows. The post-afternoon-feeding increase of H_2_ production in BS cows was not significant, averaging by 1.5 folds. The effect of 3-NOP at 1000 h led to the greatest increase in H_2_ production in both BS and HF cows, however, with the increase being more pronounced for HF compared with BS cows as opposed to CH_4_ production.

## Discussion

This study aimed to investigate the varying mitigation efficacy of 3-NOP on enteric CH_4_ emissions in HF and BS cows, based on previously observed discrepancies between these two breeds (Ma et al., 2024). Additionally, we aimed to study the potential combined effects of 3-NOP and TAN. It is important to note that both HF and BS cows were kept under the same management regimen on the research farm since their first lactation prior to the experiment. The consistency in housing and feeding conditions minimizes the influence of differing management practices, allowing us to focus on the study’s objectives related to potential breed differences.

### Milk production

In this study, we observed that 3-NOP tended to increase milk yield and ECM in HF, whereas it decreased in BS cows. This is in line with a year-long study by van Gastelen et al. (2024), who observed an increase in ECM by 6.5% with 3-NOP-supplemented HF cows. Similarly, we observed that 3-NOP tended to increase milk fat yield in both breeds and increased milk lactose yield only in HF cows. It was reported that cows on a late-lactation diet supplemented with 3-NOP had higher milk lactose content during experimental weeks 1 to 4 compared to those without 3-NOP (van Gastelen et al., 2024). The increase in milk fat yield observed in the present study aligns with findings from Lopes et al. (2016), Melgar et al. (2020a; 2021), and van Gastelen et al. (2024). Furthermore, long-term studies, including those by Melgar et al. (2020a; 2021) and van Gastelen et al. (2024), suggested that 3-NOP supplementation may enhance milk fat concentration and yield by altering rumen fermentation patterns and increasing the availability of fatty acid precursors (e.g., butyrate). In contrast, no effect was observed on milk production and composition in previous findings with 3-NOP using corn-silage or grass-silage-based diets by Hristov et al. (2015), van Gastelen et al. (2022), and recently, by Ma et al. (2024) supplemented with 3-NOP and whole cottonseed. The varying effects of 3-NOP supplementation on lactational performance in dairy cows may be attributed to the differences in experimental design, study duration, and individual animal variabilities, and further confirmatory research is required. It is important to note that while short-term crossover design studies may not fully capture treatment responses that develop over time, particularly those related to altered nutrient partitioning, most lactation-related responses tend to be consistent across Latin Square and block designs, where crossover setups closely resemble continuous studies (Huhtanen and Hetta, 2012; Zanton, 2019).

In contrast to the observed effects of 3-NOP, TAN supplementation did not affect milk yield or ECM in the current study. Similar to our findings, Lazzari et al. (2023) reported that diets with *Acacia mearnsii* extract at 2% of DM did not affect milk yield while reducing DMI by 1.30 kg/d. In line with our findings, they observed a 1.40 kg/d reduction in ECM, indicating that the tannin extracts might influence the nutritional efficiency of the diet, leading to less energy being available for milk production and milk component yield (Lazzari et al., 2023). Further, TAN supplementation reduced MUN concentration by 18.6% in the current study, indicating decreased N availability in the rumen, consistent with findings that tannins reduce ruminal protein degradation (Dschaak et al., 2011; Gerlach et al., 2018; Aguerre et al., 2020). Additionally, we noted a tendency for decreased milk true protein yield by 2.68% with TAN supplementation. Similarly, Gerlach et al. (2018) noted a reduction in milk protein yield, linked to diminished AA delivery to the duodenum, while Lazzari et al. (2023) reported reductions in milk protein concentration and yield with diets including *Acacia mearnsii*, also affecting ECM negatively. However, Aguerre et al. (2016; 2020) reported an increase in milk true protein concentration with quebracho–chestnut tannin extract at 1.80% of DM, though milk protein yield declined with higher tannin levels. Altogether, these findings (Benchaar et al., 2008; Dschaak et al., 2011; Schmithausen et al., 2018) reflect those in the current study, indicating that overall dietary tannins had little to no impact on milk component concentrations or yields, while notably reducing excessive N secretion into milk as MUN.

### Nutrient intake, apparent total-tract digestibility, and N partitioning

The results highlight breed differences, with HF cows consuming 3.7 kg/d more DMI than BS cows, enabling greater milk production and improved feed efficiency. In contrast, DMI was unaffected by 3-NOP supplementation in our study, consistent with previous findings, including a recent experiment from our group (Ma et al., 2024), a palatability study by Melgar et al. (2020b) across various 3-NOP doses, and a meta-analysis by Kim et al. (2019). However, other studies have reported that 3-NOP reduces DMI in dairy cows (Kjeldsen et al., 2024; Maigaard et al., 2024). The potential negative effect of 3-NOP on DMI may be related to increased propionate formation in the rumen that induces satiety signaling in dairy cows according to hepatic oxidation regulation (Allen, 2014; 2020). In contrast, TAN supplementation resulted in a 1 kg/d reduction in DMI, though it did not affect milk yield or ECM. In the current study, a tendency of 3-NOP and TAN interaction suggests 3-NOP may influence the reduction in DMI caused by TAN, aligning with evidence that 3-NOP generally has minimal or no impact on DMI while TAN alone had a greater impact on reduced DMI. The impact of TAN on DMI has been variable; *Acacia mearnsii* at 2% of DM generally reduced DMI (Grainger et al., 2009; Kozloski et al., 2012; Lazzari et al., 2023), while it remained unchanged with *Acacia mearnsii* at 1 to 3% of DM in other studies (Orlandi et al., 2015; Gerlach et al., 2018) and increased at 4.1% of DM in sheep (Carulla et al., 2005).

In this experiment, the ATTD of OM and ADF, remained unaffected by 3-NOP, consistent with findings from previous studies (Hristov et al., 2015; Ma et al., 2024). The results indicate breed-specific effects of TAN and 3-NOP on NDF digestibility, with HF cows showing a greater reduction (26.5%) compared to BS cows (4.47%), likely due to differences in microbial populations and their fiber-degrading capacity. Feeding TAN may exacerbate this reduction by inhibiting cellulolytic bacteria, complexing with lignocellulose, or both, thereby reducing fiber digestion (McSweeney et al., 2001). Similarly, the 3-NOP × Breed interaction highlighted contrasting effects; 3-NOP tended to increase NDF digestibility in BS cows but decreased in HF cows. This aligns with the complex role of H_2_ dynamics in rumen fermentation, where elevated H_2_ concentrations (under methanogenesis inhibited by 3-NOP) can inhibit fiber degradation (Leng, 2014; Ma et al., 2019). However, previous studies have shown that 3-NOP often has minimal effects on NDF digestion (Reynolds et al., 2014; Jayanegara et al., 2018), likely due to the utilization of alternative H_2_ sinks such as butyrate and lactate (van Lingen et al., 2016). The TAN supplementation in the current study reduced CP digestibility, likely due to tannin’s well-known effect by binding with dietary protein, thereby decreasing rumen degradation and leading to lower MUN and UUN concentrations (Aguerre et al., 2016; 2020). This binding may also inhibit enzymatic activity in the intestine, potentially reducing protein absorption (Silanikove et al., 1994; Waghorn, 2008).

The tendency for a combined effect of 3-NOP and TAN supplementation on N utilization in dairy cows indicated a greater reduction in urinary N (g/d) excretion by 7.63% when TAN was fed alone and 6.11% while fed a combination of 3-NOP and TAN. Further, a TAN × Breed interaction revealed that urinary N reductions were greater in BS (25.8%) compared to HF cows (20.2%), potentially reflecting breed-specific responses. Notably, cows fed TAN at 3% of DM exhibited a 23.0% reduction in urinary N and a 36.7% increase in fecal N in the current study, indicating a shift in N excretion from urine to feces which minimizes volatile N losses (Ndegwa et al., 2008; Ahnert et al., 2015; Orlandi et al., 2015). This shift can reduce NH_3_ and N_2_O emissions, with the decrease in urinary N resulting from a reduced ruminal NH_3_ formation due to tannin-protein complexes, and the increase in fecal N stemming from post-ruminal complexes and tannin binding in the small intestine (Eckard et al., 2010; Fagundes et al., 2021).

### Combined effect of 3-NOP and TAN on enteric gas emissions

One objective of the current study was to assess the combined effects of 3-NOP and TAN on CH_4_ mitigation in terms of the mitigation efficacy of 3-NOP. Contrary to expectations of a synergistic or additive effect, we did not observe any interactions between 3-NOP and TAN for CH_4_ production and intensity. The lack of interaction between 3-NOP and TAN was partly attributed to the overall insignificant effect of TAN supplementation on methane parameters. However, in HF cows, CH_4_ yield decreased by 24.9% when fed 3-NOP alone, suggesting a breed-specific effect in reducing enteric CH_4_ yield. In contrast, BS cows exhibited a modest reduction (4.74%) while fed 3-NOP regardless of TAN supplementation. Breed-specific differences in rumen microbial composition and fermentation dynamics likely influenced the efficacy of TAN and 3-NOP, as further discussed in the next section regarding the interaction of 3-NOP and breed.

The effect of TAN on CH_4_ mitigation has varied across studies. Schmithausen et al. (2018) found no impact of *Acacia mearnsii* at 3% of DM on CH_4_ emissions, which aligns with our findings. Conversely, Grainger et al. (2009) and Lazzari et al. (2023) reported CH_4_ reductions of 13-16% and 10%, respectively, with *Acacia mearnsii* extract at 1.8% and 2% of DM. Lazzari et al. (2023) attributed this reduction to decreased fiber degradation and reduced H_2_ availability for methanogens, though no significant changes were observed in CH_4_ yield or intensity (g/kg ECM). Additionally, Alves et al. (2017) reported a 30% decrease in CH_4_ intensity when *Acacia mearnsii* extract at 2% of DM was used with concentrate in dairy cows grazing tropical pearl millet (*Pennisetum glaucum* L) pasture. Similarly, Denninger et al. (2020) reported a reduction in CH_4_ production and yield by 18 and 16%, respectively, using *Acacia mearnsii* at 3% of DM. To balance CH_4_ mitigation with maintaining DMI and production performance, we also selected a 3% DM dose of TAN for this study, aiming to achieve effective CH_4_ reduction without compromising animal performance.

In this study, HF cows supplemented with 3-NOP showed a 22% decrease in CH_4_ production, aligning with previous findings but less than the 30% reduction reported by Dijkstra et al. (2018) and Kebreab et al. (2023). A year-long study with Holstein cows by van Gastelen et al. (2024) showed a 21, 20, and 27% reduction in CH_4_ production, yield, and intensity, respectively. The effectiveness of 3-NOP varies with diet type, and duration of supplementation and is negatively affected by high dietary NDF content. For this experiment, with a dietary NDF of 41% of DM, predictions using a model from Kebreab et al. (2023) estimated CH_4_ reductions of 26.3, 26.0, and 24.7% of CH_4_ production, yield, and intensity compared with the actual reductions of 22.0, 21.5, and 23.5% observed in HF cows. Despite the high NDF content, these results are within the expected efficacy range for 3-NOP for HF cows whereas in BS cows the reduction was lower, which was observed by Ma et al. (2024). Although Ma et al. (2024) and the current study were conducted in the same research farm, a distinct batch of individual cows were enrolled in the current study to avoid potential confounding. The 3-NOP × Breed interactions related to CH_4_ emissions are discussed further below.

Recent studies have explored combined dietary strategies to enhance the efficacy of 3-NOP in reducing CH_4_ emissions. For instance, Maigaard et al. (2024) reported that combining fat, nitrate, and 3-NOP did not benefit CH_4_ reductions greater than those achieved with 3-NOP alone. Further, Gruninger et al. (2022) reported that while 3-NOP and canola oil individually reduced enteric CH_4_ yield by 28.2% and 24.0%, respectively, their combined use resulted in an additive 51.3% reduction. However, Kjeldsen et al. (2024) observed no synergistic effects between 3-NOP and cracked rapeseed on CH_4_ production, yield, or intensity. Likewise, our recent study indicated that combining 3-NOP with whole cottonseed did not enhance the CH_4_ mitigation efficacy of 3-NOP in a high-NDF (43% of DM) diet (Ma et al., 2024). These findings suggest that the benefits of combinations depend on the specific additives used, the dietary context, and their eventual impact on microbial activity and fermentation pathways in the rumen.

Previous studies have consistently reported an increase in H_2_ emissions following CH_4_ reduction due to 3-NOP supplementation (Hristov et al., 2015; van Gastelen et al., 2022; Ma et al., 2024), with our findings confirming a 2.58-fold increase in BS cows and a 3.71-fold increase in HF cows. The rise in H_2_ emissions corresponds with the lowest CH_4_ emissions and is linked to inhibited methanogenesis causing H_2_ accumulation in the rumen (Trei et al., 1972; Janssen, 2010). Furthermore, changes in fermentation patterns and VFA proportions influence H_2_ emissions when 3-NOP is used (Reynolds et al., 2014; Lopes et al., 2016; Melgar et al., 2020a). The differences in H_2_ emissions between breeds may partly be explained by the variation in DMI. With higher DMI in HF cows than BS cows, they likely experience more intense rumen fermentation conditions due to the greater availability of fermentable substrates in the rumen, leading to stimulate higher microbial activity (Münger and Kreuzer, 2006; Rooke et al., 2014). Additionally, microbial activity likely plays a key role in shaping rumen fermentation dynamics and CH_4_ formation, as the methanogen abundance has been shown to differ between these breeds (Gonzalez-Recio et al., 2018). Contrarily, daily CO_2_ emissions remained unchanged by 3-NOP and TAN, aligning with previous findings (Hristov et al., 2015; Lopes et al., 2016), except for a reported 3% increase by Ma et al. (2024). However, similar to our findings, van Gastelen et al. (2022) reported an increase in CO_2_ yield could result from CH_4_ inhibition and a shift towards propionate, highlighting the complex impact of 3-NOP on ruminal fermentation dynamics and energy retention. This illustrates how both 3-NOP and TAN differently influence enteric gas emissions in dairy cows.

### Interaction of breed and 3-NOP on enteric gas emissions

To our knowledge, Ma et al. (2024) was the first to demonstrate the interaction between 3-NOP and dairy cattle breed on enteric CH_4_ mitigation, reporting significant interactions on CH_4_ production (HF: 18 vs. BS: 8%) and intensity (HF: 19 vs. BS: 4%), but not on CH_4_ yield (HF: 18 vs. BS: 10%). In our study, cows were also fed a high-NDF basal diet. The CH_4_ mitigation (g/d) efficacy in HF cows in both studies was consistent with the meta-analysis by Kebreab et al. (2023), which focused mostly on HF cows. However, the efficacy in BS cows was lower in both studies, indicating a breed-specific effect of 3-NOP on absolute CH_4_ emissions. In the current study, a 3-NOP × TAN × Breed interaction was observed for CH_4_ yield, which seemed to be mainly driven by the 3-NOP × Breed 2-way interaction. The 3-NOP × Breed interaction was significant for absolute CH_4_, whereas a tendency was shown for CH_4_ yield and intensity. However, it is important to note that all three CH_4_ emission metrics had similar magnitude of change between the two breeds. This outcome may be attributed to the sample size of 8 cows per breed, which was determined based on power analysis for absolute CH_4_ production according to the study objective. This could potentially affect the inference of other variables such as CH_4_ yield and intensity, especially with added variabilities from DMI and ECM.

The CH_4_ mitigation efficacy differences between breeds could be partly attributed to differences in DMI, with BS cows consuming nearly 3.7 kg/d (15% of daily intake) less than HF cows. This intake disparity likely leads to changes in rumen retention time, buffering capacity, and carbohydrate fermentation, ultimately influencing ruminal methanogenesis (Allen, 1996; Russell and Rychlik, 2001; Münger and Kreuzer, 2006) as the rumen microbial community varies with diet, time after feeding, and genotype (Rooke et al., 2014). Specifically, the varying mitigation efficacy of 3-NOP in the two breeds could be linked to rumen microbial activities, affected by feed intake (Söllinger et al., 2018). Although feeding behavior patterns, especially the conditioned meals (immediately following the fresh feed delivery) did not seem to differ between the two breeds, distinct responses in H_2_ and CH_4_ productions were observed associated with afternoon feeding. This pattern aligns with the findings by Ma et al. (2024) who noted that the effectiveness of 3-NOP decreased by evening, with the lowest emissions occurring 1 to 2 h post-afternoon feeding. A direct association was observed between FI patterns and CH_4_ emissions, where peak FI after morning feeding corresponded with increased CH_4_ production, consistent with studies by Niu et al. (2014) and DeVries et al. (2003). This underlines the strong link between diurnal feeding behaviors and CH_4_ emission patterns, further highlighted by Hristov and Melgar (2020) who observed a peak reduction of enteric CH_4_ emission by 45.2% with 3-NOP supplementation with once-a-day feeding coinciding with high FI shortly after morning feeding. Overall, these observations underscore the efficacy of 3-NOP supplementation in reducing enteric CH_4_ emissions and illustrate its dependency on feeding time dynamics.

Moreover, the above-mentioned breed-specific effects could be confounded by both host genetics and microbial activities (Difford et al., 2018). Previous studies have shown that microbial community structure and activity contribute to the CH_4_-emitting phenotype (Shi et al., 2014; Roehe et al., 2016). *Methanobrevibacter*, more prevalent in HF cows (Gonzalez-Recio et al., 2018), is more sensitive to 3-NOP, suggesting that HF cows may require lower dosages, whereas *Methanosphaera*, requiring higher doses (Pitta et al., 2022), suggesting differential effects on individual methanogenic lineages and may contribute to breed-specific responses. Additionally, inhibition of *Methanobrevibacter* by 3-NOP could shift microbial populations through microbe-microbe interactions (Henderson et al., 2015). Recently, Li et al. (2024) reported breed-specific CH_4_ yield differences, with the HF rumen microbiome favoring methylotrophic methanogenesis, reducing reliance on H_2_ + CO_2_ pathways. In contrast, Jersey cows utilize H_2_ through the Wood–Ljungdahl pathway, showing distinct microbial adaptations in H_2_ processing between breeds. However, breed-specific differences in rumen fermentation or microbial communities cannot be confirmed in this study, and future investigations should integrate host genetic and microbial factors to better understand feed efficiency and enteric CH_4_ emission interactions. In particular, studies with larger and similar experimental units from different farms and regions should be considered to further evaluate these differences.

Under methanogenesis inhibition, alternative H_2_ pathways, such as propionate, butyrate, lactate, and ethanol biosynthesis, are promoted (Mackie et al., 2023; Ungerfeld and Pitta, 2024). Acetate, butyrate, and H_2_ production are directly linked to microbial activities, while CH_4_ and propionate production are thermodynamically favored under high H_2_ conditions (Wolin et al., 1997; Janssen, 2010; Ungerfeld and Pitta, 2024). In our study, HF cows showed a greater increase in H_2_ production (271%) compared with BS cows (157%) when 3-NOP was supplemented. To assess H_2_ balance during methanogenesis-inhibited conditions, we calculated the CH_4_ reduction between CON and 3-NOP-supplemented cows, using the stoichiometric relationship of CH_4_ production from CO_2_ and H_2_ as the basis for the calculation (Thauer et al., 1998). The reductions of CH_4_ were 106 g/d for HF cows and 55 g/d for BS cows, corresponding to 53.0 g/d and 27.5 g/d of H_2_ not used for methanogenesis, respectively. These values are much higher than the H_2_ measured from eructation, which was 3.28 g/d for HF cows and 2.26 g/d for BS cows. The fate of excess H_2_ in this context remains uncertain from stoichiometric calculations alone. This uncertainty suggests that excess H_2_ may be redirected to alternative metabolic pathways. As proposed by Li et al. (2024), distinct microbial H_2_ and reductant disposal pathways may drive interbreed variations in CH_4_ emissions. However, to fully understand the metabolic fate of the unaccounted-for H_2_ under inhibition of methanogenesis, further research is needed to fully explore the mechanisms involved in this metabolic redirection.

The recent approval of 3-NOP by EFSA (2021) and FDA (2024) reinforces its role in reducing CH_4_ emissions in dairy cattle. However, both our study and Ma et al. (2024) highlight potential breed-specific variations in its efficacy, potentially influenced by feed intake, and rumen microbiome, although further investigations are needed to assess the responses on larger populations, potentially also expending to other breeds. These findings underscore the need for more comprehensive studies to optimize CH_4_ mitigation strategies and ensure their integration into GHG inventories and carbon footprint models.

## Conclusion

Data from this study showed no synergistic or additive effects on enteric CH_4_ mitigation when 3-NOP and TAN were combined, likely due to the effect of TAN remaining insignificant. However, an interaction between 3-NOP and Breed on absolute CH_4_ production was observed, while HF cows were more responsive to 3-NOP (22%) compared with BS cows (13%). Enteric CH_4_ yield and intensity also showed tendencies for 3-NOP mitigation efficacies across these two breeds. Such differences may stem from varying DMI and microbial fermentation between breeds. Supplementation of TAN reduced urinary N excretion by 23% and increased fecal N by 37%. Overall, the results highlight the importance of considering breed differences when developing and implementing CH_4_ mitigation strategies, as well as their implications for national GHG inventories. Further research is needed to evaluate the interactions between dairy cattle breeds and enteric CH_4_ mitigation strategies and to better understand their modes of action.

### Ethics approval

All animal procedures were reviewed and approved by the Cantonal Veterinary Office of Zürich, Switzerland (license no. ZH207/2021).

## Data and model availability statement

The data supporting the findings of this study are not deposited in a public repository but are available from the corresponding author upon request.

## Declaration of generative AI and AI-assisted technologies in the writing process

The author(s) did not use any AI or AI-assisted technologies in the preparation of the original manuscript.

## Acknowledgments

We extend our heartfelt appreciation to the dedicated barn staff led by Mirjam Klöppel (Strickhof, Lindau, Switzerland) and the research team led by Dr. Melissa Terranova (AgroVet-Strickhof, Lindau, Switzerland). We extend special thanks to Raphael Jendly (ETH Zürich, Switzerland) and Ueli Buchmann (Strickhof, Lindau, Switzerland) for their technical expertise with the GreenFeed system. Additionally, we appreciate Dr. Usman Arshad (ETH Zürich, Switzerland) for his insightful suggestions during the review process.

