## Supplementary Table S1 to S7 for "Divergent effects of 3-nitrooxypropanol on enteric methane emissions in Holstein and Brown Swiss cows, and its lack of synergy with *Acacia mearnsii* tannin extract"

*animal* journal

**Supplementary Table S1**

Lactational performance of Brown Swiss (BS) and Holstein Friesian (HF) cows fed dietary treatments^1^.

|  | Breed | | | | | | | | | | |  | *P*-value^2^ | | | | | | |
| --- | --- | --- | --- | --- | --- | --- | --- | --- | --- | --- | --- | --- | --- | --- | --- | --- | --- | --- | --- |
| Items | BS | | | | |  | HF | | | | |  |  |  |  |  |  |  |  |
|  | CON | 3-NOP | TAN | 3-NOP + TAN | SE^3^ |  | CON | 3-NOP | TAN | 3-NOP  +  TAN | SE |  | 3-NOP | TAN | Breed | 3-NOP ×  TAN | 3-NOP × Breed | TAN ×  Breed | 3-NOP ×  TAN  ×  Breed |
| DMI^4^, kg/d | 21.2 | 21.1 | 19.6 | 20.1 | 0.668 |  | 24.9 | 24.4 | 23.8 | 24.1 | 0.676 |  | 0.87 | 0.003 | <0.001 | 0.14 | 0.55 | 0.25 | 0.97 |
| Yield, kg/d |  |  |  |  |  |  |  |  |  |  |  |  |  |  |  |  |  |  |  |
| Milk^5^ | 29.5 | 29.3 | 29.3 | 28.4 | 1.56 |  | 38.5 | 39.7 | 37.8 | 38.8 | 1.55 |  | 0.53 | 0.17 | <0.001 | 0.60 | 0.13 | 0.75 | 0.79 |
| ECM^6^ | 28.9 | 28.4 | 29.5 | 27.6 | 1.71 |  | 37.4 | 39.4 | 37.1 | 37.8 | 1.71 |  | 0.93 | 0.49 | <0.001 | 0.28 | 0.12 | 0.55 | 0.96 |
| Milk fat^7^ | 1.06 | 1.07 | 1.10 | 1.02 | 0.723 |  | 1.39 | 1.47 | 1.43 | 1.41 | 0.072 |  | 0.94 | 0.86 | <0.001 | 0.11 | 0.35 | 0.87 | 0.97 |
| Milk true  Protein^8^ | 0.95 | 0.94 | 0.97 | 0.96 | 0.062 |  | 1.27 | 1.29 | 1.22 | 1.26 | 0.063 |  | 0.73 | 0.061 | <0.001 | 0.41 | 0.46 | 0.85 | 0.56 |
| Milk lactose^9^ | 1.40 | 1.40 | 1.38 | 1.35 | 0.075 |  | 1.85 | 1.93 | 1.81 | 1.86 | 0.743 |  | 0.19 | 0.062 | <0.001 | 0.42 | 0.083 | 0.68 | 0.89 |
| Milk composition, % | | |  |  |  |  |  |  |  |  |  |  |  |  |  |  |  |  |  |
| Milk fat | 3.59 | 3.67 | 3.82 | 3.60 | 0.141 |  | 3.68 | 3.64 | 3.82 | 3.71 | 0.142 |  | 0.25 | 0.22 | 0.68 | 0.15 | 0.99 | 0.83 | 0.42 |
| Milk true  protein | 3.30 | 3.29 | 3.35 | 3.46 | 0.092 |  | 3.33 | 3.41 | 3.25 | 3.33 | 0.092 |  | 0.83 | 0.054 | 0.49 | 0.98 | 0.54 | 0.95 | 0.29 |
| Milk lactose | 4.71 | 4.72 | 4.73 | 4.75 | 0.032 |  | 4.73 | 4.77 | 4.76 | 4.79 | 0.032 |  | 0.088 | 0.21 | 0.11 | 0.97 | 0.53 | 0.92 | 0.93 |
| MUN, mg/dL | 17.3 | 16.5 | 13.5 | 14.1 | 0.465 |  | 16.7 | 16.1 | 13.5 | 12.6 | 0.488 |  | 0.24 | <0.001 | 0.096 | 0.48 | 0.45 | 0.70 | 0.22 |
| SCC, ×10^3^ cells/mL | 117 | 118 | 116 | 117 | 0.271 |  | 118 | 122 | 117 | 114 | 0.270 |  | 0.47 | 0.45 | 0.62 | 0.21 | 0.49 | 0.28 | 0.39 |
| ECM:DMI | 1.31 | 1.37 | 1.45 | 1.38 | 0.810 |  | 1.52 | 1.60 | 1.53 | 1.57 | 0.081 |  | 0.42 | 0.41 | 0.016 | 0.25 | 0.51 | 0.23 | 0.59 |

^1^Dietary treatments were CON = basal diet; 3-NOP = 3-nitrooxypropanol supplemented diet (60 mg/kg DM); TAN = Acacia mearnsii tannin extract supplemented diet (3% of DM; extract containing 51.8% total tannin/kg of DM; CT:HT ratio was 0.219); 3-NOP + TAN supplemented diet. [3-NOP No] is the LSM of CON and TAN diet; [3-NOP Yes] is the LSM of 3-NOP and 3-NOP + TAN diet; [TAN No] is the LSM of CON and 3-NOP diet; [TAN Yes] is the LSM of TAN and 3-NOP + TAN diet.

^2^*P*-values derived from the comprehensive model.

^3^Reported the largest SE.

^4^DMI, 3-NOP × TAN interaction; [3-NOP No TAN No] = 23.0^a^, [3-NOP Yes TAN No] = 22.7^ab^, [3-NOP No TAN Yes] = 21.7^c^, [3-NOP Yes TAN Yes] = 22.0^bc^. Different superscripts indicate a significant (*P* < 0.05) difference.

^5^Milk production, 3-NOP × Breed interaction; [BS 3-NOP No] = 29.3^b^, [BS 3-NOP Yes] = 28.8^b^, [HF 3-NOP No] = 38.1^a^, [HF 3-NOP Yes] = 39.2^a^. Different superscripts indicate a significant (*P* < 0.05) difference.

^6^ECM, 3-NOP × Breed interaction; [BS 3-NOP No] = 29.0^b^, [BS 3-NOP Yes] = 28.0^b^, [HF 3-NOP No] = 37.3^a^, [HF 3-NOP Yes] = 38.8^a^. Different superscripts indicate a significant (*P* < 0.05) difference. ECM yield (kg/d) = 12.18 × Fat [kg/d] + 7.69 × Protein [kg/d] + 5.26 × Lactose [kg/d] + 0.01 × Milk Yield [kg/d] (Sjaunja et al., 1990).

^7^Fat yield, 3-NOP × TAN interaction; [3-NOP No TAN No] = 1.22^a^, [3-NOP Yes TAN No] = 1.27^a^, [3-NOP No TAN Yes] = 1.26^a^, [3-NOP Yes TAN Yes] = 1.22^a^. Different superscripts indicate a significant (*P* < 0.05) difference.

^8^Milk true protein = milk crude protein (%) - MUN (mg/dL) × 6.38 / 1000.

^9^Lactose yield, 3-NOP × Breed interaction; [BS 3-NOP No] = 1.39^b^, [BS 3-NOP Yes] = 1.38^b^, [HF 3-NOP No] = 37.3^a^, [HF 3-NOP Yes] = 38.8^a^. Different superscripts indicate a significant (*P* < 0.05) difference.

**Supplementary Table S2**

Digestible nutrients^1^ and apparent total-tract digestibility (ATTD)^2^ of Brown Swiss (BS) and Holstein Frisian (HF) cows fed dietary treatments^3^.

|  | Breed | | | | | | | | | | |  | *P*-value^4^ | | | | | | |
| --- | --- | --- | --- | --- | --- | --- | --- | --- | --- | --- | --- | --- | --- | --- | --- | --- | --- | --- | --- |
| Apparent digestibility, % | BS | | | | |  | HF | | | | |  |  |  |  |  |  |  |  |
|  | CON | 3-NOP | TAN | 3-NO  + TAN | SE^5^ |  | CON | 3-NOP | TAN | 3-NOP + TAN | SE |  | 3-NOP | TAN | Breed | 3-NOP ×  TAN | 3-NOP × Breed | TAN ×  Breed | 3-NOP ×  TAN  ×  Breed |
| OM | 62.7 | 65.3 | 65.1 | 66.3 | 2.11 |  | 63.7 | 65.1 | 66.0 | 65.1 | 2.10 |  | 0.34 | 0.25 | 0.89 | 0.40 | 0.56 | 0.79 | 0.83 |
| NDF^6^ | 35.2 | 54.3 | 45.2 | 45.8 | 4.63 |  | 52.8 | 46.3 | 38.0 | 37.9 | 4.63 |  | 0.33 | 0.15 | 0.71 | 0.36 | 0.12 | 0.067 | 0.20 |
| ADF | 43.1 | 44.6 | 40.2 | 40.4 | 3.16 |  | 46.4 | 41.9 | 41.2 | 42.4 | 3.16 |  | 0.79 | 0.11 | 0.59 | 0.51 | 0.55 | 0.72 | 0.35 |
| Starch^7^ | 99.3 | 99.3 | 99.1 | 99.2 | 0.134 |  | 99.4 | 99.5 | 99.4 | 99.7 | 0.146 |  | 0.015 | 0.90 | 0.080 | 0.13 | 0.25 | 0.064 | 0.45 |
| CP | 78.7 | 79.1 | 76.4 | 76.3 | 1.73 |  | 80.2 | 79.4 | 79.4 | 78.3 | 1.73 |  | 0.78 | 0.080 | 0.050 | 0.74 | 0.65 | 0.45 | 0.92 |

^1^Digestible nutrient = nutrient intake (kg/d) × ATTD (%) of nutrient.

^2^Apparent total-tract digestibility (ATTD, %) = 100 – 100 × [(% marker in feed ÷ % marker in feces) × (% nutrient in feces ÷ % nutrient in feed)] based on Keulen and Young (1977).

^3^Dietary treatments were CON = basal diet; 3-NOP = 3-nitrooxypropanol supplemented diet (60 mg/kg DM); TAN = *Acacia mearnsii* tannin extract supplemented diet (3% of DM; extract containing 51.8% total tannin/kg of DM; CT:HT ratio was 0.219); 3-NOP + TAN supplemented diet. [3-NOP No] is the LSM of CON and TAN diet; [3-NOP Yes] is the LSM of 3-NOP and 3-NOP+ TAN diet; [TAN No] is the LSM of CON and 3-NOP diet; [TAN Yes] is the LSM of TAN and 3-NOP+ TAN diet.

^4^*P*-values derived from the comprehensive model.

^5^Reported the largest SE.

^6^NDF, TAN × Breed interaction; [BS TAN No] = 44.7^a^, [BS TAN Yes] = 42.7^a^, [HF TAN No] = 49.5^a^, [HF TAN Yes] = 36.4^a^. 3-NOP × Breed interaction; [BS 3-NOP No] = 37.5^a^, [BS 3-NOP Yes] = 49.1^a^, [HF 3-NOP No] = 44.2^a^, [HF 3-NOP Yes] = 42.6^a^. Different superscripts indicate a significant (*P* < 0.05) difference.

^7^Starch, TAN × Breed interaction; [BS TAN No] = 99.3^a^, [BS TAN Yes] = 99.2^a^, [HF TAN No] = 99.5^a^, [HF TAN Yes] = 99.5^a^. 3-NOP × TAN interaction; [3-NOP No TAN No] = 99.4^a^, [3-NOP Yes TAN No] = 99.4^a^, [3-NOP No TAN Yes] = 99.3^a^, [3-NOP Yes TAN Yes] = 99.4^a^. Different superscripts indicate a significant (*P* < 0.05) difference.

**Supplementary Table S3**

Nitrogen utilization and purine derivatives (PD) excretion in Brown Swiss (BS) and Holstein Friesian (HF) cows fed dietary treatments^1^.

|  | Breed | | | | | | | | | | |  | *P*-value^2^ | | | | | | |
| --- | --- | --- | --- | --- | --- | --- | --- | --- | --- | --- | --- | --- | --- | --- | --- | --- | --- | --- | --- |
| Items | BS | | | | |  | HF | | | | |  |  |  |  |  |  |  |  |
|  | CON | 3-NOP | TAN | 3-NOP +  TAN | SE^3^ |  | CON | 3-NOP | TAN | 3-NOP +  TAN | SE |  | 3-NOP | TAN | Breed | 3-NOP × TAN | 3-NOP × Breed | TAN ×  Breed | 3-NOP ×  TAN  ×  Breed |
| N intake, g/d | 577 | 565 | 500 | 530 | 35.1 |  | 621 | 680 | 633 | 635 | 35.0 |  | 0.28 | 0.077 | <0.001 | 0.85 | 0.63 | 0.28 | 0.22 |
| N excretion or secretion, g/d | | | | | |  |  |  |  |  |  |  |  |  |  |  |  |  |  |
| Urinary N^4^ | 130 | 129 | 121 | 122 | 1.56 |  | 131 | 132 | 121 | 123 | 1.56 |  | 0.37 | <0.001 | 0.29 | 0.14 | 0.61 | 0.42 | 0.73 |
| Urinary urea-N | 42.3 | 43.9 | 31.4 | 30.7 | 3.39 |  | 45.7 | 47.7 | 31.5 | 35.0 | 3.40 |  | 0.31 | <0.001 | 0.29 | 0.90 | 0.54 | 0.63 | 0.58 |
| Fecal N | 160 | 170 | 249 | 253 | 20.7 |  | 176 | 171 | 282 | 263 | 20.7 |  | 0.84 | <0.001 | 0.17 | 0.64 | 0.48 | 0.55 | 0.87 |
| Total excreta N | 290 | 299 | 370 | 375 | 21.0 |  | 307 | 303 | 403 | 386 | 20.9 |  | 0.89 | <0.001 | 0.15 | 0.71 | 0.51 | 0.60 | 0.85 |
| Milk N | 170 | 165 | 150 | 157 | 9.46 |  | 210 | 227 | 203 | 212 | 9.43 |  | 0.49 | 0.006 | <0.001 | 0.78 | 0.17 | 0.65 | 0.26 |
| N excretion and  secretion | 459 | 457 | 486 | 523 | 24.8 |  | 519 | 529 | 606 | 602 | 24.8 |  | 0.84 | <0.001 | <0.001 | 0.91 | 0.97 | 0.73 | 0.69 |
| N partition, % of daily intake | | |  |  |  |  |  |  |  |  |  |  |  |  |  |  |  |  |  |
| Urinary N^5^ | 23.4 | 23.1 | 17.4 | 17.3 | 0.337 |  | 23.0 | 22.7 | 18.2 | 18.2 | 0.337 |  | 0.47 | <0.001 | 0.35 | 0.55 | 0.92 | 0.013 | 0.63 |
| Urinary urea-N | 7.67 | 7.56 | 6.43 | 5.67 | 0.692 |  | 7.31 | 6.66 | 4.85 | 5.13 | 0.691 |  | 0.39 | <0.001 | 0.024 | 0.85 | 0.78 | 0.56 | 0.32 |
| Fecal N | 28.2 | 30.6 | 49.7 | 48.5 | 5.11 |  | 28.5 | 25.3 | 44.8 | 41.9 | 5.09 |  | 0.65 | <0.001 | 0.13 | 0.75 | 0.59 | 0.53 | 0.68 |
| Total excreta N | 51.2 | 53.4 | 74.0 | 71.8 | 6.05 |  | 49.7 | 44.7 | 64.2 | 61.5 | 6.04 |  | 0.54 | <0.001 | 0.021 | 0.86 | 0.63 | 0.44 | 0.67 |
| Milk N | 30.2 | 27.8 | 30.5 | 29.9 | 2.47 |  | 33.7 | 33.6 | 32.4 | 33.8 | 2.47 |  | 0.73 | 0.79 | 0.046 | 0.47 | 0.45 | 0.46 | 0.98 |
| N excretion and  secretion | 82.0 | 80.6 | 97.9 | 94.7 | 6.87 |  | 83.1 | 78.9 | 96.0 | 96.9 | 6.86 |  | 0.28 | <0.001 | 0.56 | 0.74 | 0.61 | 0.63 | 0.37 |
| Urine output, kg/d | 26.4 | 25.9 | 21.9 | 22.9 | 1.73 |  | 30.0 | 31.3 | 26.9 | 30.0 | 1.73 |  | 0.13 | 0.002 | 0.001 | 0.30 | 0.31 | 0.32 | 0.92 |
| Urinary PD excretion, mM/d | | | | |  |  |  |  |  |  |  |  |  |  |  |  |  |  |  |
| Uric acid^6^ | 86.6 | 68.8 | 76.0 | 65.4 | 9.77 |  | 86.2 | 114 | 80.8 | 91.6 | 9.76 |  | 0.62 | 0.070 | <0.001 | 0.64 | 0.014 | 0.50 | 0.29 |
| Allantoin | 815 | 810 | 787 | 802 | 104.1 |  | 971 | 1015 | 940 | 931 | 103.8 |  | 0.84 | 0.53 | 0.011 | 0.88 | 0.93 | 0.71 | 0.76 |
| Total PD | 902 | 887 | 857 | 868 | 106.5 |  | 1058 | 1128 | 1021 | 1018 | 106.3 |  | 0.77 | 0.39 | 0.006 | 0.83 | 0.80 | 0.71 | 0.69 |

^1^Dietary treatments were CON = basal diet; 3-NOP = 3-nitrooxypropanol supplemented diet (60 mg/kg DM); TAN = *Acacia mearnsii* tannin extract supplemented diet (3% of DM; extract containing 51.8 g total tannin/kg of DM and CT:HT = 0.219); 3-NOP + TAN supplemented diet. [3-NOP No] is the LSM of CON and TAN diet; [3-NOP Yes] is the LSM of 3-NOP and 3-NOP+ TAN diet; [TAN No] is the LSM of CON and 3-NOP diet; [TAN Yes] is the LSM of TAN and 3-NOP+ TAN diet.

^2^*P*-values derived from the comprehensive model.

^3^Reported the largest SE.

^4^Urinary N excretion (g/d), 3-NOP × TAN interaction; [3-NOP No TAN No] = 131^a^, [3-NOP Yes TAN No] = 130^a^, [3-NOP No TAN Yes] = 121^b^, [3-NOP Yes TAN Yes] = 123^b^. Different superscripts indicate a significant (*P* < 0.05) difference.

^5^Urinary N (% of daily intake), TAN × Breed interaction; [BS TAN No] = 23.3^a^, [BS TAN Yes] = 17.3^b^, [HF TAN No] = 22.8^a^, [HF TAN Yes] = 18.2^b^. Different superscripts indicate a significant (*P* < 0.05) difference.

^6^Uric acid excretion (mmol/d), 3-NOP × Breed interaction; [BS 3-NOP No] = 81.3^bc^, [BS 3-NOP Yes] = 69.4^c^, [HF 3-NOP No] = 85.8^ab^, [HF 3-NOP Yes] = 99.5^a^. Different superscripts indicate a significant (*P* < 0.05) difference.

|  | Breed | | | | | | | | | | |  | *P*-value^2^ | | | | | | |
| --- | --- | --- | --- | --- | --- | --- | --- | --- | --- | --- | --- | --- | --- | --- | --- | --- | --- | --- | --- |
| Items | BS | | | | |  | HF | | | | |  |  |  |  |  |  |  |  |
|  | CON | 3-NOP | TAN | 3-NOP +  TAN | SE^3^ |  | CON | 3-NOP | TAN | 3-NOP  + TAN | SE |  | 3-NOP | TAN | Breed | 3-NOP × TAN | 3-NOP × Breed | TAN ×  Breed | 3-NOP ×  TAN  ×  Breed |
| CH_4_ production^4^, g/d | 422 | 376 | 430 | 362 | 23.5 |  | 517 | 390 | 469 | 379 | 23.6 |  | <0.001 | 0.19 | 0.045 | 0.73 | 0.050 | 0.20 | 0.22 |
| CH_4_ yield^5^, g/kg DMI | 19.0 | 18.1 | 21.9 | 18.3 | 1.21 |  | 20.9 | 15.7 | 19.4 | 15.8 | 1.21 |  | <0.001 | 0.54 | 0.10 | 0.63 | 0.13 | 0.066 | 0.10 |
| CH_4_ intensity^6^, g/kg MY | 14.4 | 12.4 | 15.4 | 13.0 | 0.879 |  | 13.6 | 9.90 | 12.5 | 10.1 | 0.875 |  | <0.001 | 0.69 | <0.001 | 0.51 | 0.35 | 0.12 | 0.32 |
| CH_4_ intensity^7^, g/kg ECM | 14.7 | 12.8 | 14.9 | 13.4 | 0.881 |  | 13.9 | 10.1 | 12.6 | 10.2 | 0.878 |  | <0.001 | 0.91 | <0.001 | 0.23 | 0.15 | 0.19 | 0.61 |
| H_2_ production, g/d | 1.76 | 3.55 | 1.22 | 3.75 | 0.663 |  | 1.09 | 4.47 | 1.22 | 4.58 | 0.661 |  | <0.001 | 0.74 | 0.33 | 0.41 | 0.23 | 0.48 | 0.45 |
| H_2_ yield, g/kg DMI | 0.077 | 0.174 | 0.047 | 0.189 | 0.029 |  | 0.046 | 0.180 | 0.051 | 0.191 | 0.029 |  | <0.001 | 0.99 | 0.76 | 0.40 | 0.63 | 0.62 | 0.57 |
| CO_2_ production^8^, g/d | 12804 | 12349 | 11463 | 12738 | 410 |  | 14122 | 14261 | 13765 | 13976 | 410 |  | 0.15 | 0.087 | <0.001 | 0.032 | 0.64 | 0.70 | 0.075 |
| CO_2_ yield, g/kg DMI | 575 | 597 | 601 | 662 | 30.7 |  | 577 | 577 | 569 | 595 | 30.6 |  | 0.090 | 0.16 | 0.074 | 0.32 | 0.48 | 0.21 | 0.85 |
| O_2_ consumption, g/d | 8675 | 8652 | 8563 | 8955 | 501 |  | 9170 | 9290 | 9262 | 9365 | 501 |  | 0.54 | 0.74 | 0.12 | 0.68 | 0.90 | 0.98 | 0.69 |

**Supplementary Table S4**

Enteric gas emissions of Brown Swiss (BS) and Holstein Friesian (HF) cows fed dietary treatments^1^.

^1^Dietary treatments were CON = basal diet; 3-NOP = 3-nitrooxypropanol supplemented diet (60 mg/kg DM); TAN = *Acacia mearnsii* tannin extract supplemented diet (3% of DM; extract containing 51.8 g total tannin/kg of DM and CT:HT = 0.219); 3-NOP + TAN supplemented diet. [3-NOP No] is the LSM of CON and TAN diet; [3-NOP Yes] is the LSM of 3-NOP and 3-NOP + TAN diet; [TAN No] is the LSM of CON and 3-NOP diet; [TAN Yes] is the LSM of TAN and 3-NOP + TAN diet.

^2^*P*-values derived from the comprehensive model.

^3^Reported the largest SE.

^4^CH_4_ production, 3-NOP × Breed interaction; [BS 3-NOP No] = 421^b^, [BS 3-NOP Yes] = 366^c^, [HF 3-NOP No] = 489^a^, [HF 3-NOP Yes] = 383^bc^. Different superscripts indicate a significant (*P* < 0.05) difference.

^5^CH_4_ yield, 3-NOP × TAN × Breed interaction; [3-NOP No TAN No HF] = 20.9^a^, [3-NOP Yes TAN No BS] = 18.1^ab^, [3-NOP Yes TAN No HF] = 15.7^b^, [3-NOP Yes TAN Yes BS] = 18.3^ab^, [3-NOP No TAN Yes BS] = 21.9^a^, [3-NOP No TAN Yes HF] = 19.4^ab^, [3-NOP Yes TAN Yes HF] = 15.8^b^, [3-NOP No TAN No BS] = 19.0^ab^. Different superscripts indicate a significant (*P* < 0.05) difference.

^6^CH_4_ intensity (g/kg milk yield), TAN × Breed interaction; [BS TAN No] = 13.4^ab^, [BS TAN Yes] = 14.0^a^, [HF TAN No] = 11.7^ab^, [HF TAN Yes] = 11.1^b^. Different superscripts indicate a significant (*P* < 0.05) difference.

^7^CH_4_ intensity (g/kg ECM), 3-NOP × Breed interaction; [BS 3-NOP No] = 14.7^a^, [BS 3-NOP Yes] = 13.1^a^, [HF 3-NOP No] = 13.2^a^, [HF 3-NOP Yes] = 10.1^b^. Different superscripts indicate a significant (*P* < 0.05) difference.

^8^CO_2_ production, 3-NOP × TAN × Breed interaction; [3-NOP No TAN No HF] = 14122^ab^, [3-NOP Yes TAN No BS] = 12349^d^, [3-NOP Yes TAN No HF] = 14261^a^, [3-NOP Yes TAN Yes BS] = 12738^cd^, [3-NOP No TAN Yes BS] = 11463^d^, [3-NOP No TAN Yes HF] = 13765^abc^, [3-NOP Yes TAN Yes HF] = 13976^abc^, [3-NOP No TAN No BS] = 12804^bcd^. Different superscripts indicate a significant (*P* < 0.05) difference.

**Supplementary Table S5**

Least-squares means of enteric methane emission (g/h) measured at 3-h intervals in Brown Swiss and Holstein cows.

| Items | BS | | | |  | HF | | | |
| --- | --- | --- | --- | --- | --- | --- | --- | --- | --- |
| Time of day | 3-NOP No | 3-NOP Yes | SE | % Reduction |  | 3-NOP No | 3-NOP Yes | SE | % Reduction |
| 0100 h | 16.8 | 16.7 | 0.939 | 0.595 |  | 18.9 | 16.6 | 1.19 | 12.2 |
| 0400 h | 15.4 | 13.9 | 0.939 | 9.74 |  | 16.3 | 14.5 | 1.23 | 11.0 |
| 0700 h | 13.5 | 12.8 | 0.980 | 5.19 |  | 15.1 | 14.1 | 1.19 | 6.62 |
| 1000 h | 19.5^a^ | 15.0^b^ | 0.939 | 23.1 |  | 22.1^a^ | 15.6^b^ | 1.19 | 29.4 |
| 1300 h | 19.6^a^ | 15.2^b^ | 1.03 | 22.4 |  | 22.1^a^ | 17.4^b^ | 1.23 | 21.3 |
| 1600 h | 17.3 | 14.6 | 1.02 | 15.6 |  | 20.8^a^ | 15.2^b^ | 1.19 | 26.9 |
| 1900 h | 18.6 | 16.0 | 0.939 | 14.0 |  | 22.6^a^ | 17.4^b^ | 1.19 | 23.0 |
| 2200 h | 19.1 | 16.0 | 1.03 | 16.2 |  | 21.5 | 17.0 | 1.19 | 20.9 |

**Supplementary Table S6**

Least-squares means of hydrogen emission (g/h) measured at 3-h intervals in Brown Swiss and Holstein cows.

| Items | BS | | | | |  | HF | | | | |
| --- | --- | --- | --- | --- | --- | --- | --- | --- | --- | --- | --- |
| Time of day | 3-NOP No | 3-NOP Yes | SE | Fold-increase | % Increase |  | 3-NOP No | 3-NOP Yes | SE | Fold-increase | % Increase |
| 0100 h | 0.034 | 0.149 | 0.028 | 4.38 | 338 |  | 0.046 | 0.148 | 0.030 | 3.22 | 222 |
| 0400 h | 0.039 | 0.054 | 0.028 | 1.38 | 38.5 |  | 0.038 | 0.042 | 0.030 | 1.11 | 10.5 |
| 0700 h | 0.034 | 0.082 | 0.029 | 2.41 | 141 |  | 0.039 | 0.048 | 0.030 | 1.23 | 23.1 |
| 1000 h | 0.045 | 0.212 | 0.029 | 4.71 | 371 |  | 0.060 | 0.273 | 0.029 | 4.55 | 355 |
| 1300 h | 0.063 | 0.196 | 0.031 | 3.11 | 211 |  | 0.057 | 0.229 | 0.030 | 4.02 | 302 |
| 1600 h | 0.033 | 0.086 | 0.030 | 2.61 | 161 |  | 0.030 | 0.139 | 0.030 | 4.63 | 363 |
| 1900 h | 0.062 | 0.145 | 0.028 | 2.34 | 134 |  | 0.081 | 0.249 | 0.029 | 3.07 | 207 |
| 2200 h | 0.089 | 0.153 | 0.031 | 1.72 | 71.9 |  | 0.052 | 0.220 | 0.030 | 4.23 | 323 |

**Supplementary Table S7**

Least-squares means of feed intake rate (%/h) of Brown Swiss and Holstein cows fed twice daily with or without 3-NOP supplementation.

| Items | BS | | |  | HF | | |
| --- | --- | --- | --- | --- | --- | --- | --- |
| Time of day | 3-NOP No | 3-NOP Yes | SE |  | 3-NOP No | 3-NOP Yes | SE |
| 0000h | 1.11 | 1.16 | 0.449 |  | 1.05 | 1.28 | 0.592 |
| 0100h | 1.99 | 1.57 | 0.465 |  | 2.25 | 2.04 | 0.792 |
| 0200h | 2.14 | 2.26 | 0.502 |  | 2.28 | 2.10 | 0.619 |
| 0300h | 2.21 | 2.15 | 0.503 |  | 2.25 | 1.89 | 0.619 |
| 0400h | 0.91 | 0.90 | 0.481 |  | 1.00 | 1.15 | 0.619 |
| 0500h | 3.56 | 3.78 | 0.464 |  | 3.46 | 4.51 | 0.570 |
| 0600h | 4.89 | 4.89 | 0.449 |  | 5.36 | 5.54 | 0.594 |
| 0700h | 0.46 | 1.07 | 0.455 |  | 0.56 | 1.04 | 0.795 |
| 0800h | 5.02 | 4.13 | 0.471 |  | 3.79 | 3.08 | 0.552 |
| 0900h | 10.7 | 11.0 | 0.501 |  | 12.1 | 12.4 | 0.570 |
| 1000h | 4.52 | 5.21 | 0.435 |  | 6.30 | 5.60 | 0.570 |
| 1100h | 5.39 | 5.58 | 0.435 |  | 5.64 | 5.37 | 0.570 |
| 1200h | 4.51 | 4.57 | 0.482 |  | 5.00 | 5.04 | 0.570 |
| 1300h | 4.01 | 4.97 | 0.449 |  | 4.54 | 5.07 | 0.619 |
| 1400h | 4.26 | 4.26 | 0.449 |  | 4.30 | 4.78 | 0.570 |
| 1500h | 4.46 | 3.62 | 0.449 |  | 4.00 | 5.50 | 0.570 |
| 1600h | 4.37 | 4.59 | 0.449 |  | 3.59 | 3.70 | 0.595 |
| 1700h | 8.58 | 6.86 | 0.464 |  | 6.79 | 7.26 | 0.594 |
| 1800h | 11.1 | 11.0 | 0.435 |  | 11.6 | 10.7 | 0.570 |
| 1900h | 4.40 | 5.89 | 0.435 |  | 6.13 | 4.63 | 0.592 |
| 2000h | 3.74 | 4.41 | 0.435 |  | 4.19 | 3.48 | 0.570 |
| 2100h | 3.67 | 3.66 | 0.449 |  | 3.71 | 3.69 | 0.619 |
| 2200h | 2.69 | 3.07 | 0.448 |  | 2.67 | 3.87 | 0.592 |
| 2300h | 2.54 | 2.64 | 0.483 |  | 2.39 | 3.10 | 0.594 |
